# Listening to the room: disrupting activity of dorsolateral prefrontal cortex impairs learning of room acoustics in human listeners

**DOI:** 10.1101/2025.04.04.644835

**Authors:** Heivet Hernández-Pérez, Jessica J.M. Monaghan, Jason Mikiel-Hunter, James Traer, Paul F Sowman, David McAlpine

**Author notes:** Corresponding author, Heivet Hernandez-Perez, **Email:**.

## Abstract

Navigating complex sensory environments is critical to survival, and brain mechanisms have evolved to cope with the wide range of surroundings we encounter. To determine how listeners learn the statistical properties of acoustic spaces, we assessed their ability to perceive speech in a range of noisy and reverberant rooms. Listeners were also exposed to repetitive transcranial stimulation (rTMS) to disrupt the dorsolateral prefrontal cortex (dlPFC) activity, a region believed to play a role in statistical learning. Our data suggest listeners rapidly adapt to statistical characteristics of an environment to improve speech understanding. This ability is impaired when rTMS is applied bilaterally to the dlPFC. The data demonstrate that speech understanding in noise is best when exposed to a room with reverberant characteristics common to human-built environments, with performance declining for higher and lower reverberation times, including fully anechoic (non-reverberant) environments. Our findings provide evidence for a reverberation “sweet spot” and the presence of brain mechanisms that might have evolved to cope with the acoustic characteristics of listening environments encountered every day.

## Introduction

Learning occurs over multiple time scales—evolutionary, developmental and moment-to-moment—to support a diverse range of abilities: navigating complex sensory environments (Bregman, 1994; Lewicki et al., 2014; Smith & Lewicki, 2006), expressing intricate behaviours in social settings (Gariépy et al., 2014; van den Bos et al., 2013), or acquiring communication skills such as bird song (Brainard & Doupe, 2002; Lauay et al., 2004; Marler, 1970) or spoken language (Aslin et al., 1998; Saffran et al., 1996; Saffran, 2003). Although some learning requires active or explicit involvement in a task (Huyck & Wright, 2011; Mathews et al., 1989; Rebuschat, 2015), perhaps with a system of rewards and punishments to make the learning ‘stick’ (Barberis, 2013; Schultz, 2002; Wächter et al., 2009), other forms of learning seem automatic or implicit (Reber, 1967), acquired with individuals seemingly unaware it is taking place. This type of learning, referred to as ‘statistical learning’ (Ambrus et al., 2020; Saffran et al., 1996), ‘sequence learning’ (Nissen & Bullemer, 1987; Vékony et al., 2022) or ‘sequential learning’ (Conway & Christiansen, 2001; Vékony et al., 2022), is thought to entail automatic and incidental extraction of regularities or patterns within external stimuli or in the environment (Conway, 2020; Takács et al., 2021). Evident across sensory modalities to support automatic learning of tonal (Saffran et al., 1999) or linguistic (Saffran et al., 1996) sequences, strings of letters (Reber, 1967), visual scenes and shapes (Fiser & Aslin, 2001), visual-motor patterns (Nissen & Bullemer, 1987), and even tactile input (Conway & Christiansen, 2005), statistical learning appears to be a unitary, domain-general phenomenon, potentially governed by a single mechanism or neurocognitive principle (Conway, 2020; Kirkham et al., 2002).

Increasing evidence suggests that statistical learning is a critical part of how listeners deal with complex and cluttered acoustic scenes. Potential background sounds such as rain or insects—referred to as sound textures (Hicks & McDermott, 2024; McDermott et al., 2013; McWalter & McDermott, 2018) as well as changes in the regularity of sound patterns (Barascud et al., 2016; Bianco et al., 2020) are processed—seemingly unconsciously—in terms of their summary statistics. This form of statistical learning likely contributes to our ability to follow conversations in background noise (‘cocktail party listening’; Cherry, 1953) and to deal with reverberant spaces where multiple, delayed copies of the same sound reach a listener in the form of reflections from walls and other acoustically opaque surfaces (Blesser & Salter, 2009; Sabine, 1953; Schroeder, 1962). Though listeners rely on early-arriving sound energy to determine source location—suppressing potentially conflicting localization cues in later-arriving, often more intense, sound energy (Bradley et al., 1999; Culling et al., 2003; Houtgast & Steeneken, 1985; Nielsen & Dau, 2010)—the perception of reflected sound energy is informative of the listening environment more broadly (Bronkhorst & Houtgast, 1999; Shinn-Cunningham, 2000). This includes whether environments are real or synthetic i.e., deviated from natural reverberation characteristics (Traer & McDermott, 2016), room dimensions (Brumm & Naguib, 2009; Cabrera et al., 2005; Kolarik et al., 2021; Zahorik & Wightman, 2001) and source distance (Bronkhorst & Houtgast, 1999; Zahorik & Wightman, 2001).

Nevertheless, despite its potential utility for understanding background features of a sound environment, the accumulation over time of late-arriving, reverberant sound energy is thought to generate an additional burden on listening performance beyond that from sound energy direct from interfering sources (Houtgast & Steeneken, 1973; Knudsen, 1929; Lochner & Burger, 1961; Santon, 1976; Shinn-Cunningham & Kawakyu, 2003), smearing the acoustic waveform and occluding temporal gaps that might otherwise be helpful for ‘glimpsing’ speech in background noise (Cooke, 2006). However, if listeners are able to utilize late-arriving, reverberant energy of known acoustic environments to suppress disruptive spatial cues and enhance speech understanding (Brandewie & Zahorik, 2010, 2013; Vlahou et al., 2019; Watkins, 2005a, 2005b), then the acoustic characteristics of an environment can be learned and stored for later use in complex listening tasks.

Here, using an ecologically relevant listening task—understanding speech in background noise—we assessed the ability of human listeners to learn the statistical structure of different sound environments defined by their RT_60,_ the time it takes for reverberant energy to decay by 60 decibels (dB), and confirmed that speech understanding in background noise improved over time. Specifically, when asked to report words from unfamiliar and semantically uninformative spoken sentences of varying duration in noise convolved with the reverberant qualities of different acoustic environments, listeners’ performance improved with increasing sentence duration, and with repeated exposure to each environment (Brandewie & Zahorik, 2010, 2013).

Given these findings, we wondered, if the learning of acoustic environments i.e., improvements in performance with increasing sentence duration, follow a similar time-course to *in vivo* experimental evidence of increased adaptive capacity for neural learning of sound environments upon repeated exposure to those environments (Dean et al., 2005, 2008), and whether this adaptive capacity is modulated by cortical circuits (Robinson et al., 2016) such as the dorsolateral prefrontal cortex (dlPFC), a cortical locus hypothesised to contribute to statistical learning in human listeners (Ambrus et al., 2020; Vékony et al., 2022). Specifically, we asked whether knowledge of the acoustic environment accumulated over time—that is, whether performance during initial exposures is poorer than during later ones—and whether this accumulation is reduced when bilateral dlPFC activity is disrupted by TMS. We found that the capacity to learn the acoustic environment to aid speech understanding was diminished when repetitive transcranial magnetic stimulation (rTMS) was applied bilaterally to impair the function of dlPFC, suggesting that indeed human listeners have neural mechanisms supporting listening in noise and reverberant conditions.

Consistent with our findings, Francl and McDermott (2022) demonstrated that an auditory model trained under realistic listening conditions—incorporating both noise and reverberation—was able to reproduce several human-like spatial hearing characteristics. We therefore explored whether the learning effects we observed relied on the combination of real-world environments we employed, i.e., the RT_60_s in which listening performance was assessed. Specifically, we hypothesized that the ability to understand speech in noise depends on exposure to specific values of RT_60_ across our experimental paradigm, including those in the range commonly experienced by human listeners in natural and built environments (Traer & McDermott, 2016). Counter to expectations that reverberant energy can only harm speech understanding, listeners’ abilities to leverage the knowledge of a sound environment (talker identity, speech corpus, noise characteristics, spatial configuration) to improve speech understanding was best when a ‘typical’ room with reverberant characteristics close to the average of a wide range of common (built) environments (Traer & McDermott, 2016) was included as one of the three environments presented in a single experimental run. Performance declined systematically when this room was switched for one with more, or less, reverberation, including a fully anechoic (i.e., non-reverberant or dry) environment in which listeners initially trained on the recall task. Importantly in the context of other potentially learnable acoustic features, talker identity (three male and three female) represented a random variable in our experimental design. Our evidence for a reverberation ‘sweet spot’ suggests the existence of brain mechanisms adapted to the longer-term structure of common listening environments (Traer & McDermott, 2016).

## Materials and Methods

### Participants

A total of 74 participants (53 were females), aged between 19-26 years old (mean ± SD = 22 ± 2 years old) were recruited across five experiments. All participants were naïve to the experimental paradigm. In Experiment 1, 22 participants were tested in three rooms: a Lecture Room, an Open-Plan Office, and an Underground Car Park. Experiment 2 used the same three rooms combined with transcranial magnetic stimulation (TMS), with participants receiving either sham (n = 11) or real (n = 10) stimulation. Experiments 3-5 each used the Open-Plan Office and Underground Car Park, paired with a third room that differed across experiments: an Anechoic Room (Experiment 3; n = 10), a Living Room (Experiment 4; n = 11), or a Highly Reflectant Room (Experiment 5; n = 10). All participants were Australian native-English speakers, had normal pure tone thresholds (< 20 dB HL tested at octave intervals between 0.5-8 kHz (Hughson & Westlake, 1944); Interacoustics Hearing Aid Fitting Analyzer Affinity 2.0 Audiometry) and normal middle ear function (assessed using standard 226 Hz tympanometry; Titan, Interacoustics). To ensure normal outer hair cell function, all participants were screened for Distortion Product Otoacoustic Emissions (DPOAEs) between 0.5-10 kHz (stimulus parameters: f1/f2 =1.2, f1 =65 dB SPL; f2 =55 dB SPL; responses parameters: SNR > 6 dB, signal level > −10 dB SPL, and reliability > 98% (DPOAE440; Titan, Interacoustics). All participants had normal steady-state ipsilateral broadband noise middle ear muscle reflexes (MEMR) (Titan, Interacoustics). The audiological testing took 15-20 minutes to complete.

### Acoustic stimuli

To assess listeners’ abilities to understand speech in background noise across different sound environments, they were asked to verbally report keywords spoken by a talker virtually located in front of the participant that were masked by white noise arriving from a virtual source 90° to the left (Figure 1A) (Brandewie & Zahorik, 2010, 2013). A binaural configuration was necessary as speech intelligibility improvements following exposure to room acoustics is significantly reduced under monaural listening conditions (Brandewie & Zahorik, 2010). The speech stimuli used in this study were from the Coordinate Response Measure (CRM) corpus (Bolia et al., 2000), and all combinations of Callsigns (‘Baron’, ‘Eagle’, ‘Charlie’, ‘Tiger’, ‘Arrow’, ‘Ringo’, ‘Laker’, ‘Hoper’), Colors (four monosyllabic choices, ‘red’, ‘white’, ‘blue’, or ‘green’), and Numbers (the English digits between ‘one’ and ‘eight’) were used in this experiment (Figure 1B) (Brandewie & Zahorik, 2010, 2013). The masking white noise was randomly generated on a PC using MATLAB (The MathWorks, Inc. of Natick, MA: https://au.mathworks.com/) and was presented with the speech. The masking noise preceded the speech stimuli by 150 ms, during which its amplitude increased linearly from 0 to the target level (70 dB SPL) — i.e., a 150-ms linear onset ramp. The masker offset coincided with the end of the speech stimuli and was not ramped (Brandewie & Zahorik, 2013).

**Fig 1.**
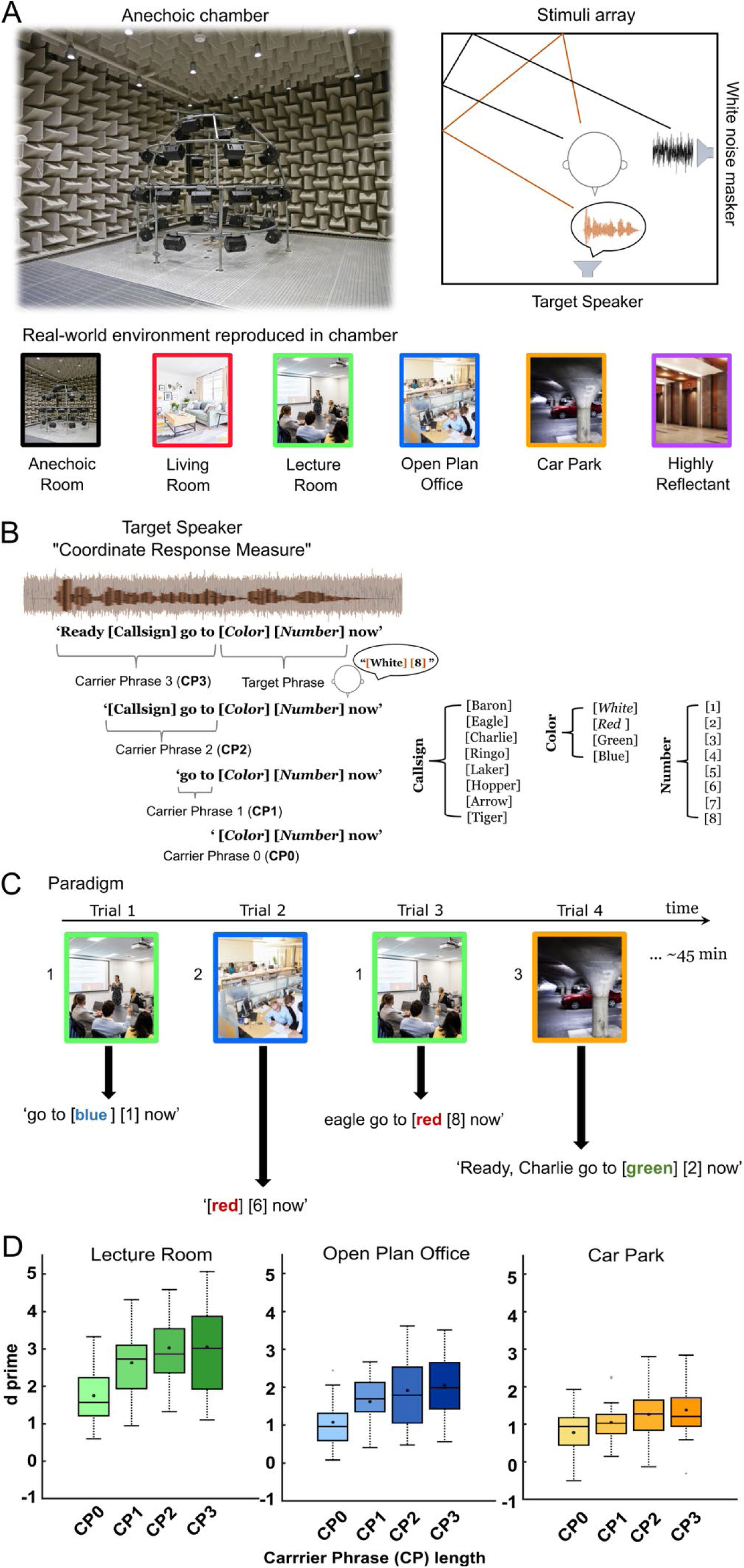
Experimental environment. **A.** Speech-in-noise immersed in real reverberant environments reproduced in an anechoic chamber over a 41-loudspeaker array. All speech material was rendered as if it was originating from directly in front of a listener and was masked by spatially separated (90° to the left) white noise at −15 dB signal-to-noise ratio (SNR). The noise masker level was fixed at 70 dB SPL. The speech material was convolved with the measured impulse response of a variety of different real rooms, from anechoic to highly reverberant real rooms such as an underground car park (see Table 1 methods for acoustic details). **B.** Structure of the speech material. Sentences from the Coordinated Response Measures (CRM) corpus were modified to vary their duration depending on how many words preceded the target phrase i.e., ‘carrier phrase’ (CP) length. Participants were asked to recognize and repeat target words: |Color| from a list of four and |Number| from a 1-8 list. **C.** Example of the experimental paradigm. Trials consisted of ‘carrier phrases’ spoken in a specific room i.e., ‘go to blue 1 now’ in the Lecture Room. Participants were never exposed to the same room consecutively and the task lasted no longer than 45 minutes. **D.** Sensitivity to |Color| and |Number| combined in the Lecture Room, Open-Plan Office and Underground Car Park, shown separately by carrier phrase (CP) length for transparency; see Methods and Results for statistical treatment of this factor. Mean d’ (measure of accuracy calculated as: Z(correct responses) – Z(false alarms)) denoted as circles in the boxplot (n=22 for all rooms). The horizontal line denotes the median. Upper and lower limits of the boxplot represent 1st (Q1) and 3rd (Q3) quartiles respectively, while whiskers denote the interquartile range (IQR = Q3–Q1),data available: https://doi.org/10.25949/24295342. Only room images in this figure were generated using artificial intelligence, with the exception of the Anechoic Room photograph, which was taken by the authors in the chamber where the experiments took place.

**Table 1.** Real rooms acoustic characteristics.

|  | **RT<sub>60iso</sub> (ISO 3382) / RT<sub>60Traer</sub> (Traer & McDermott, 2016)** | **Room Volume** | **Critical distance** | **Distance [m]** | **DRR [dB]** | **C50 [dB]** |
|---|---|---|---|---|---|---|
| Anechoic Room | 0/0 | n/a | n/a | 1.3 | n/a | n/a |
| Living room | 0.33s/0.40s | 62.47m<sup>3</sup> | 0.85m | 1.3 | 19.2 | 26.7 |
| Lecture Room | 0.46s/0.49s | 164m<sup>3</sup> | 2.78m | 1.3 | 6.87 | 15.44 |
| Open-Plan Office | 0.96s/0.85s | 446m<sup>3</sup> | 2.49m | 1.3 | 5.2 | 13.89 |
| Small Reflectant room | 1.55s/1.27s | 134m<sup>3</sup> | 1.18m | 1.3 | 0.29 | 3.83 |
| Underground Car Park | 2.42s/1.32s | >5700m<sup>3</sup> | 3.34m | 1.3 | 7.21 | 11.53 |

DRR: Direct to Reverberant Ratio
C50: Speech Clarity Index

### Virtual Environments

Reverberant environments were reproduced by convolving the generated acoustic signals with Room Impulse Responses (RIRs) obtained using 62-channel microphone array recordings of real rooms that were subsequently decoded into 41 higher-order Ambisonic (HOA) channels (Badajoz-Davila et al., 2020; Weisser et al., 2019). These final decoded channels corresponded to the spherical array of 41 Tannoy V8 concentric loudspeakers (Tannoy) installed in the anechoic chamber at the Australian Hearing Hub where testing took place. To optimize the “directionality” of the acoustic stimuli, the speech and masking noise signals were separately convolved with RIRs that had enhanced direct sound components in loudspeakers associated with their respective virtual source locations (i.e., 0° azimuth loudspeaker for the target talker and 90° azimuth loudspeaker for the masking noise) (Badajoz-Davila et al., 2020; Weisser et al., 2019). In the anechoic condition, RIRs only incorporating the enhanced direct sound components were convolved with the speech and masking noise signals. To prevent confounding changes in overall audibility across environments and carrier-phrase lengths—which would occur if only the direct-sound components were equalized across sources with differing Direct-to-Reverberant Energy Ratios (DRRs), (see Table 1)—stimuli were normalized by their overall broadband RMS level (direct plus reverberant energy) at the listener position using an omnidirectional reference microphone at the array centre (Badajoz-Davila et al., 2020; Kressner et al., 2018; Weisser et al., 2019). During data collection, the SNR was fixed at −15 dB after RIR convolution by adjusting the target speech gain (measured over the target phrase) relative to a fixed convolved masker level of 70 dB SPL. Holding overall broadband levels constant across environments attenuated the direct-sound component in each reverberant room relative to the anechoic condition according to each source’s DRR (Badajoz-Davila et al., 2020; Kressner et al., 2018; Weisser et al., 2019).

All signal processing was performed on a PC using MATLAB (Mathworks) and the final spatialized signal were presented via the RME MADI sound card (RME Audio) to two RME 32-channel digital-to-analog converters (M-32, RME Audio). These, in turn, fed 11 Yamaha XM4180 power amplifiers (Yamaha) that drove the loudspeaker array.

### Identity of sound environments

Three rooms were selected, with similar Reverberation Times (RT_60_s) to those employed by Brandewie and Zahorik (2013) and assessed in 22 participants. As Bajadoz-Davila et al. (2020) and Brandewie and Zahorik (2013) calculated these RT_60_s according to ISO-3382 guidelines (ISO, 2009), they are referred to in the text as RT_60iso_: Lecture room (RT_60iso_ = 0.46 s), Open-Plan Office (RT_60iso_ = 0.96 s) and Underground Car Park (RT_60iso_ = 2.42 s) (Table 1). Additionally, we ran three follow-up experiments in separate groups of naïve listeners, each pairing the Open-Plan Office and Underground Car Park with a different third room (Anechoic Room, Living Room, or Highly Reflectant Room). These conditions were employed to determine whether a specific combination was required to observe improvements in speech performance as exposure to room acoustics increased, that is, with increasing ‘carrier phrase length’, assessed in the same way as (Brandewie & Zahorik, 2013). For each combination, Lecture Room was swapped with one of three virtual environments: Anechoic (RT_60iso_ = n/a), Living Room (RT_60iso_ = 0.33 s) and Highly Reflectant Room (RT_60iso_ of 1.55 s) (Table 1). Naïve participants were recruited for each room combination.

To compare our recordings to RIR data recorded and analyzed by Traer and McDermott (2016), we also calculated RT_60_s for each room by integrating across the 41 HOA channels of each RIR and then calculating the median RT_60_ value in 31 frequency sub-bands with centre frequencies ranging from 80 Hz to 10 kHz (Traer & McDermott, 2016). We refer to these measures of reverberation as RT_60traer_. The range of sub-bands and their frequency selectivity was chosen to match that of the human ear (Glasberg & Moore, 1990; McDermott & Simoncelli, 2011). See *SI Materials and Methods,* (Traer & McDermott, 2016), for further information concerning how individual RT_60traer_s were extracted from each sub-band.

### Stimulus Normalization and Binaural Rendering

Stimuli were generated and normalized within a 41-channel sound field prior to binaural rendering to preserve natural head-related acoustic effects (e.g., head shadow). Source speech and noise signals were first convolved with 41-channel anechoic or reverberant impulse responses. The overall energy of the 41-channel noise field was scaled to a target of 70 dB, and the 41-channel speech field was subsequently scaled to achieve the target SNR. Following sound field normalization, the 41-channel signals were rendered to two channels using a Higher Order Ambisonics to Binaural (hoa2bin) decoder. To verify the at-ear acoustic levels, simulated at-ear root-mean-square (RMS) energy was extracted for the speech and noise tokens post-rendering. These digital RMS values were mapped to approximate dB SPL, based on the 70 dB noise sound field anchor, to confirm the resulting at-ear acoustic SNRs across both anechoic and reverberant conditions (Table 2).

**Table 2.**
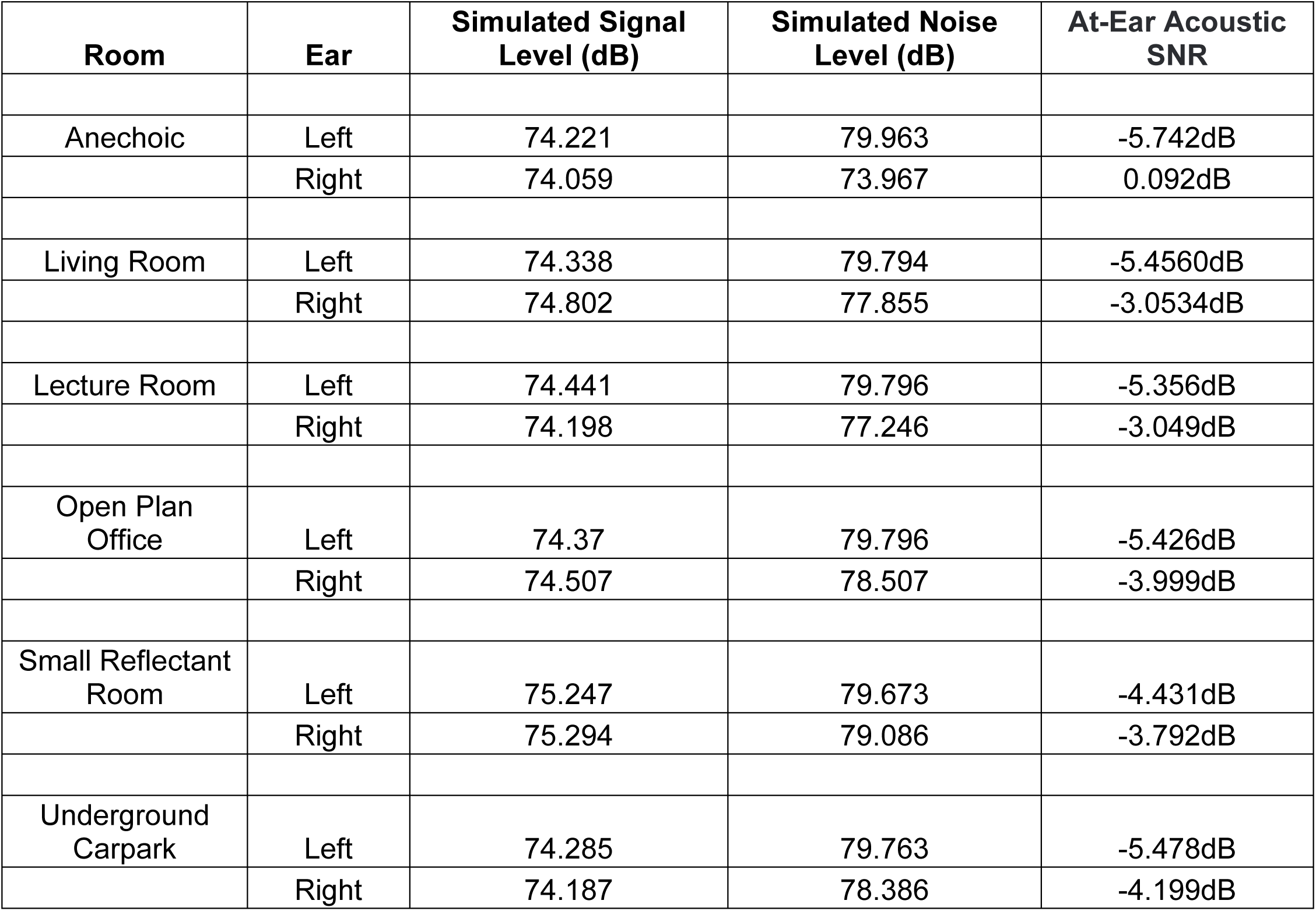
Summary at ear-acoustic levels for main rooms.

### Continuous theta-burst stimulation

Applying repetitive Transcranial Magnetic Stimulation (rTMS)—specifically continuous theta burst stimulation (cTBS)—to the right and left dlPFC, elicits a period of depressed cortical excitability in the targeted area that lasts about 60 minutes post-stimulation (Gamboa et al., 2010; Hoogendam et al., 2010; Huang et al., 2005; Romero et al., 2022), the time during which speech recall in reverberant rooms was assessed. Participants were screened for contra-indications to rTMS (see Supplemental Information. Transcranial Magnetic Stimulation screening form). Two cTBS conditions (‘real’ and ‘sham’ TMS) were counterbalanced across normal-hearing participants, and participants were blinded to the type of manipulation they might receive.

cTBS intensity applied to dlPFC was adjusted for every participant (i.e., both ‘real’ and ‘sham’ TMS groups) as a function of their individual corticospinal excitability assessed with single-pulse TMS-motor thresholds. Single-pulse TMS was delivered using a 70 mm figure-of-eight coil (Magstim Rapid2 system, Magstim, Whitland, United Kingdom), orientated at 45° to the scalp with current flowing posterior-anterior across the primary motor cortex. For the determination of single pulse TMS-induced motor thresholds we used visual observation of the first dorsal interosseous (FDI) muscle twitch (Varnava et al., 2011). Coil position and angle was adjusted until the optimal site for consistent and larger elicitation of FDI muscle twitch. Once the optimal site was located, the minimum level of stimulator output that produces a visible twitch in either the left or right hand in each individual, following single pulse stimulation of the respective contralateral motor cortex, was determined. Subjects were instructed to relax their arm, and rest it on their lap, palm upwards, whilst gently squeezing their thumb and forefinger together – as if squeezing a pea, whilst simultaneously keeping their other fingers relaxed. We then monitored for 3/5 visible twitches in the FDI in response to single pulse TMS. Subjects were occasionally asked to shake their arm to relax everything and then to resume holding their hand as before (Coltheart et al., 2018; Dienes & Hutton, 2013). Participants’ motor thresholds ranged from 46-57% of the maximum stimulator output (mean = 51 ± 5). Participants’ individual motor thresholds for left and right hemisphere were recorded to set the intensity for bilateral cTBS stimulation which was administered by placing the same figure-of-eight coil over the right and left dlPFC located over electrode positions F3 and F4 (10-20 system EEG) (Herwig et al., 2003; Jurcak et al., 2005). Bilateral TMS was performed serially i.e., first the right or the left dlFPC (order was alternated among subjects), was chosen to limit possible compensation effects of the non-stimulated hemisphere (Ambrus et al., 2020). Each cTBS burst consisted of three pulses at 50 Hz, with bursts repeated at a frequency of 5 Hz, applied continuously for 40s, and delivered at an intensity of 80% of the right or left resting motor threshold (Huang et al., 2005).

Both real (applied to 10 naïve listeners) and sham (applied to 11 naïve listeners) TMS stimulations were identical and delivered to two groups of listeners, with participants counterbalanced between the two conditions. The sole difference between the ‘real’ and ‘sham’ TMS was the figure-of-8 coil utilized. Real TMS was administered using an active figure-of-8 coil, while the sham stimulation employed a sham coil (Magstim Rapid2 system, Magstim, Whitland, United Kingdom). The sham coil is non-stimulating, meaning it does not induce a significant magnetic field, and is designed to be indistinguishable from the active coil in appearance, feel, and sound. The duration of the TMS procedure was 20–30 minutes, primarily due to the estimation of motor thresholds, with cTBS stimulation delivered in the final 2–5 minutes.

### Procedure

#### Familiarization

All participants first completed (within 5 minutes) a familiarization task that consisted of 10 trials (i.e., different ‘carrier phrase lengths’) presented in anechoic conditions at 0 dB SNR where listeners were asked to report |Color| and |Number|. Participants exposed to ‘real’ or ‘sham’ TMS completed the familiarization and behavioural task right after cTBS procedures. All participants performed the familiarization phase with 80-100% accuracy, confirming that all participants including exposed to cTBS understood the task and procedural learning was achieved and not affected by either ‘sham’ or ‘real’ TMS exposure.

#### Behavioural task

The CRM sentences were spoken by six talkers (3 males and 3 females) and were of varying duration where all, some, or none of the preceding sentence (the ‘carrier phrase’ (CP, Figure 1B)) before ‘|Color| |Number|’ was included (Brandewie & Zahorik, 2010, 2013). Thus, for CP0, listeners heard ‘|Color| |Number| now’, CP1—‘go to |Color| |Number| now’, CP2—‘|Callsign| go to |Color| |Number| now’, CP3—‘Ready |Callsign| go to |Color| |Number| now’ (Figure 1B and C). After each phrase was presented, participants reported the |Color| and |Number| they heard to the experimenter and performance was assessed based on keywords correctly identified. The rationale for only reporting |Color| and |Number| was to ensure the same amount of keywords identifications across different phrases lengths. For the reverberant conditions SNR was set to −15 dB, which we found in pilot testing resulted in a level of performance similar to (Brandewie & Zahorik, 2010, 2011, 2013; Zahorik, 2009). The room environment was selected randomly for each trial except that the same room could not appear in two consecutive trials. The target |Color|, |Number| and talker were selected randomly for each trial. For each experimental session, three different listening environments were assessed, and listeners were presented 360 trials (30 repeats x ‘room = 3’ x ‘carrier phrase length =4’). Each listener participated in one session only. Throughout the session, listeners were required to verbally repeat the appropriate |Color| and |Number| combination they heard from corpus lists physically displayed on both sides of the 0° azimuth loudspeaker. The CRM is a closed-set corpus, therefore participants were provided with |Color| and |Number| choices to select from (Brandewie & Zahorik, 2010, 2011, 2013; Zahorik, 2009). The experimenter continuously monitored participants’ responses while scoring performance, but no feedback was provided. The behavioural total test time was 45 min.

### Speech performance analysis and time course-fittings of mean cumulative hit rates

Sensitivity was estimated separately for colour and number responses in each room and carrier phrase length. For each participant, *d’* was calculated for each individual |Color| and |Number| using Equation 1, where z(H) and z(F) are the z transforms of the hit rate and false-alarm rate, respectively. A hit was a trial on which the presented |Color| or |Number| was correctly selected. A false alarm was a trial on which any incorrect |Color| or |Number| was reported I.e., the false-alarm rate for each item was averaged over the other items. To prevent infinite values of *d’*, hit or false-alarm rates of 1 were set to 0.99, and rates of 0 were set to 0.01.

We estimated sensitivity for each item because response accuracy in the CRM corpus, and confusions between colours and numbers, vary across stimuli pairs (Brungart, 2001). Our design also produced unequal target frequencies across conditions and participants. Therefore, an item-level estimate keeps these differences visible, whereas a single index over all alternatives approach (Hacker & Ratcliff, 1979) would assume equal confusability across alternatives.

Item-level *d’* values were averaged within each dimension to give a single colour *d’* and a single number *d’* for each participant, room and carrier phrase. The total *d’* for a carrier phrase in a given room was the mean of the colour and number *d’* values for that combination, and these totals were then averaged across participants.

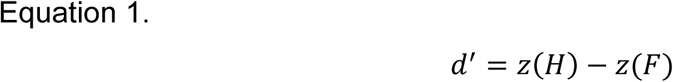

Because *d’* is calculated independently for |Color| and |Number|, it lacked the temporal resolution necessary to describe performance over time due to data paucity. Consequently, we analysed the development of individual and average performance in each acoustic environment using cumulative hit rates. We applied a 5-point moving average (∼7 seconds) to each trace and plotted performance as a function of mean cumulative exposure time.

To quantify performance trajectories, we fitted the time-series data with both single (Eq. 2) and double (Eq. 3) exponential functions:

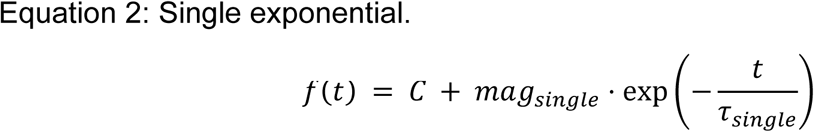

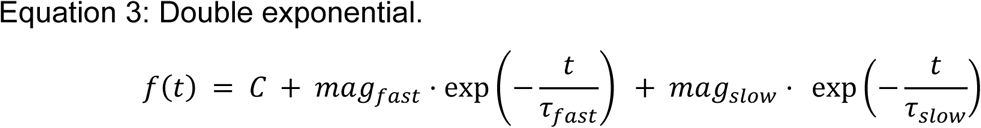

where, *f*(*t*) represents the fitted performance at time *t*, and *C* denotes the constant y-intercept. The terms *mag* and *τ* represent the magnitude and time constant (in seconds), respectively, for the single, fast, and slow components.

Constrained optimization was performed using MATLAB’s fmincon function to minimize the mean squared error. To avoid local minima, fits were initialized using a gridded search. For single exponential models, we tested a 25 × 25 grid with *τ_sinssle_* ranging from 0–100 s and *mag_sinssle_* from −50 to 50. For double exponential models, we utilized a grid of 5 linearly spaced values for *τ_sloss_* (50–500 s), *τ_fast_* (0–100 s), and their corresponding magnitudes (−10 to 10), resulting in 625 initialization points per model type. Upper bounds were set at 2000 s for *τ_sinssle_* and *τ_sloss_*, and 500 s for *τ_fast_*. The optimal fit for a listening environment was selected from the resulting collection of single/double exponential curves by identifying fits with the highest adjusted R^2^. During optimization, fits were weighted by their variance; therefore, greater importance was attributed to performance at later exposure times. Furthermore, we forced all final fitted curves to pass through the final hit rate i.e. last data point, by penalizing those that did not during fitting.

Double exponential models were fitted exclusively to the full dataset, as they are capable of accounting for early data reversals. However, to account for inconsistent initial results, single exponential models were calculated for both the full dataset and a truncated dataset (excluding the first 10 points, (∼14 seconds). To determine the optimal model, we calculated the Akaike Information Criterion (AIC). AIC estimates prediction error by balancing goodness-of-fit against model complexity, defined as:

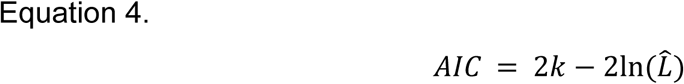

Where *k* is the number of estimated parameters and *L*^⏢^ is the maximum likelihood of the model. In the context of least-squares fitting, this is derived from the sum of squared errors. To ensure valid comparisons, AIC values were computed using identical data ranges for both candidate models (comparing fits on the full dataset against one another or fits on the truncated dataset against one another).

For analysis of ‘Learning Settling Time”, we calculated the absolute percentage deviation of cumulative hit rate curves relative to the final time point in each curve i.e., their Final Hit Rate (FHR), we then calculated back in time to determine where fitted curves first deviated from the FHR by ±10%. Note that FHR is the final data point of the cumulative hit rate i.e., after all responses have been accumulated in that listening environment.

### Statistical analysis

Repeated measures ANOVAs (rANOVA) were performed to assess speech understanding (d’) within subjects with factors: Rooms (levels: Lecture Room, Open-Plan Office and Underground Car park); carrier phrase length (levels: CP0, CP1, CP2 and CP3) and target word (levels: |Color| and |Number|). A mixed ANOVA was employed to assess whether speech understanding was affected by the between factor: TMS exposure (levels ‘sham’ and ‘real’ TMS), and within-subject’s factors: Rooms (levels: Lecture Room, Open-Plan Office and Underground Car park); carrier phrase length (levels: CP0, CP1, CP2 and CP3) and target word (levels: |Color| and |Number|). Additionally, three follow up experiments were performed and a one-way ANOVA was used to assess whether speech understanding was affected by the room-context (i.e., the 3^rd^ room in which Open-Plan Office and Underground Car Park were learned). Only one between-subject’s factor was assessed: room context.

Effect sizes were calculated for all *post hoc* statistical analyses (Partial Eta-squared (*η_p_^2^*), Cohen’s d (parametric analysis) and Person’s r (for nonparametric Wilcoxon Signed-Rank test) (Pallant, 2011). Further two-tailed t-tests (original alpha=0.05, with *post hoc* Bonferroni corrections for multiple comparisons) were performed. Corrected alpha value is not reported as all statistical analyses were performed in SPSS: RRID:SCR_002865 (IBM Corp. Released 2023. IBM SPSS Statistics for Windows, Version 29.0.2.0 Armonk, NY), and SPSS output adjusts the p-value instead of the alpha value itself.

### Power Analysis

Initial sample size estimation for individual ANOVA analysis (18) was computed using G*Power: *RRID:SCR_013726* (Faul et al., 2007) (Effect size f = 0.25; α err prob = 0.05; Power (1-β err prob) = 0.95). However, given the large effect size observed in previous studies (Brandewie & Zahorik, 2010, 2013; Srinivasan & Zahorik, 2012; Pavel Zahorik & Brandewie, 2016) using sample sizes = 9-16 listeners, we expected relatively large effect sizes with a sample ≥ 10. The variance of all variables was tested for normal distribution (Shapiro-Wilk test, Shapiro & Wilk, 1965).

### Data availability statement

All data and related metadata were deposited in Figshare, an appropriate public repository (https://doi.org/10.25949/24295342)

### Code Information

All custom code used for data collection and analysis will be available upon request.

## Results

Human listeners can incorporate knowledge of a listening environment, specifically its reverberant characteristics, to improve speech understanding in noise (Brandewie & Zahorik, 2010, 2013; Srinivasan & Zahorik, 2012; Zahorik & Brandewie, 2016). To understand how this learning of sound environments is achieved, we recreated acoustic characteristics of real rooms using an array of loudspeakers located in an anechoic chamber (Figure 1A) and assessed listeners’ performance in a speech-in-noise task using sentences from the Coordinate Response Measure (CRM) corpus—“Ready |Callsign| go to |Color| |Number| now” (Figure 1B). The CRM *corpus* is commonly used to quantify listening performance in noisy or cluttered environments (Bolia et al., 2000). We employed six environments, (Figure 1A), that varied from fully anechoic to a highly reverberant underground car park with RT_60iso_ = 2.42 s (RT_60_s according to ISO-3382 guidelines (ISO, 2009).

In our task, we asked listeners to report the |Color| (one of four monosyllabic choices, ‘red’, ‘white’, ‘blue’, or ‘green’) and the |Number| (the English digits between ‘one’ and ‘eight’) they heard for sentences of varying duration where all, some, or none of the preceding sentence (the ‘carrier phrase length’) before ‘|Color| |Number|’ was included (Figure 1B). Importantly, words in the CRM phrase preceding ‘|Color| |Number|’ are uninformative as to the color and number (Brandewie & Zahorik, 2010, 2013). CRM sentences—complete or partial (see Methods and Figure 1B)—were presented as if originating from in front of a participant, whilst a randomly generated white noise was presented from a source 90° to their left. Listeners’ abilities to recognize |Color| and |Number| from whole or partial CRM sentences were assessed. Although sentences varied in duration from trial to trial, they always contained the element ‘|Color| |Number| now’ embedded in noise and convolved with the impulse response of the real rooms (Figure 1C). Sentence length varied by adjusting the duration of the preceding ‘carrier phrase’ (CP; Figure 1B-C), which was always constructed from the same CRM sentence embedded in noise and convolved with the same impulse response as the remainder of the phrase. Thus, for CP0, listeners heard only ‘|Color| |Number| now’, CP1—‘go to |Color| |Number| now’, CP2—‘|Callsign| go to |Color| |Number| now’, CP3—‘Ready |Callsign| go to |Color| |Number| now’.

After each phrase was presented, participants verbally reported the |Color| and |Number| they heard to the experimenter, and performance was assessed based on keywords correctly identified. CRM phrases of varying length were spoken by one of 6 talkers (3 female, 3 male) and the different environments were presented in a pseudorandom order to avoid the same environment being presented in consecutive trials (e.g., Figure 1C). Our initial assessment, (Figure 1D), examined listening performance in three environments: a Lecture Room (RT_60iso_ = 0.45 s), an Open-Plan Office (RT_60iso_ = 0.96 s), and an Underground Car Park (RT_60iso_ = 2.42 s). We confirmed, in 22 listeners, that understanding speech in background noise depends on the reverberant characteristics of the listening spaces from which impulse responses were obtained [Figure 1D (all environments and CP lengths) and Figure 2A (rANOVA relevant comparisons between environments)]. Overall, listeners performed better (quantified in terms of d’ for reporting the correct |Color| and |Number|) in the Lecture Room—the least reverberant of the 3 environments (Figure 1D and 2A)—with a significant main effect of room [rANOVA: F (2,42) = 76.75, p<0.001, *η_p_^2^* = 0.79]. Bonferroni-corrected *post-hoc* pairwise comparisons demonstrated that performance in the Lecture Room was significantly better than in the Open-Plan Office [mean difference = 0.41, p < 0.001] and Underground Car Park [mean difference = 1.05, p < 0.001], (Figure 2A). Performance was also significantly better in the Open-Plan Office than in the Underground Car Park [mean difference = 0.65, p < 0.001], Figure 2A.

**Figure 2.**
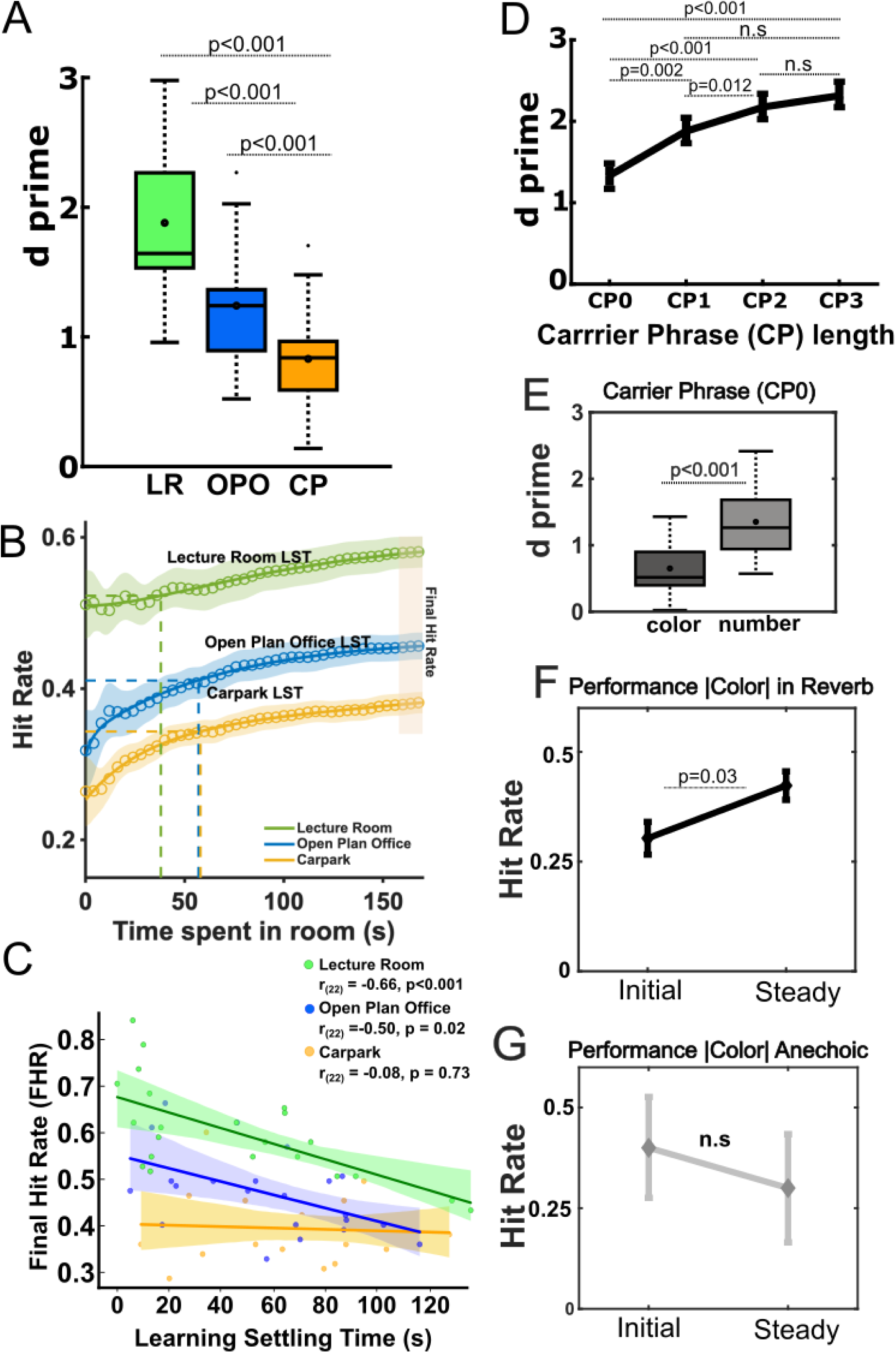
Learning effects in three real rooms. Data shown here reflects the significant effects identified in the rANOVA (see Methods); see Fig. 1d for the full dataset across all conditions. **A.** Performance (d’) (including correct and incorrect responses to all |*Colors|* and |*Numbers|*) in the Lecture Room (LR), Open-Plan Office (OPO) and Car Park (CP). Performance includes all speakers and phrases durations. Mean d’ denoted as circles in the boxplot (n=22 for all rooms). The horizontal line denotes the median. Upper and lower limits of the boxplot represent ^1s^t (q1) and ^3r^d (q3) quartiles respectively, while whiskers denote the interquartile range (IQR =q3-q1). **B.** Time course of performance in each room for 22 listeners. Correct responses (Hit Rate) across time spent in each room are shown, solid curves in color correspond to mean data after 5-time point moving averages, equivalent to ∼7s, were applied). Optimal time course-fittings (using a two-phase association model) are plotted as color markers in each room. Associated shaded areas representing standard errors of the mean. Dashed coloured lines represent the time point at which participants reached a stable performance in each environment [i.e., ± 10% of the Final Hit Rate (FHR)], here ‘Learning Settling Time (LST)’, this variable was computed for each listener. Notice that no statistical differences were found for LST among the different rooms. **C.** Shows a correlation analysis between FHR and SLT in each environment. Significant negative Pearson correlations were observed for Lecture Room, and Open Plan Office, suggesting the earlier participants reach a stable performance, the higher their FHR is. **D.** Performance for each carrier phrase length, d’ was significantly better as the ‘carrier phrases length’ increased except for CP2 vs. CP3 (n.s), where a roll over effect was observed. **E.** Performance to |Color| and |Number| for CP0 i.e., our proxy for short-term adaptation the carrier phrase where exposure to an environment remained minimal. Here all the environments have been collapsed i.e., rANOVA, main effect of “target word”. **F-G.** Hit Rate for Initial (1-2) and Steady (9-10) trials for keyword |Color| in the CP0 condition: (F) in Reverberant conditions, indicating a significant improvement in performance for Steady trials, likely due to the accumulation of acoustic/environmental knowledge i.e., meta-adaptation (G) in anechoic (no echoes) condition, showing a lack of improvement in performance i.e., lack of meta-adaptation in the absence of reverberation. All data available: https://doi.org/10.25949/24295342

### Statistical learning of reverberant environments occurs over long and short-time courses

Statistical learning likely occurs over different time courses subject to a range of possible brain mechanisms controlling or modulating the different cadences over which learning emerges (Robinson et al., 2016; Simpson et al., 2014). We sought to distinguish short-term from longer-term learning by assessing how prior exposure to the listening environment benefits word recall over multiple time courses. The design of our paradigm—with talker, length of carrier phrase, target words |Color| & |Number| and reverberation time (listening environment) constituting random variables—means that the initial phase of learning may also be highly variable; listeners, idiosyncratically, likely experience very different parameters over the first few trials, making it difficult to assess the rate at which learning accumulates over these early epochs, and we indeed found this to be the case (see Supplemental Figure 1).

To characterize behavioural response trajectories, we fitted performance data with single and double exponential functions. Double exponentials were fitted to the full time course (Supp. Fig 1 and 2). Single exponentials were fitted to both the full time course (Supp. Fig. 1) and a truncated version excluding the first 10 points (∼14 s) to mitigate initial variability in exposure to the listening configuration (Supp. Fig. 2). AIC (Akaike Information Criterion) comparisons heavily favoured the double exponential model for the full time course (mean ΔAIC Double Vs Single (fitted to full time course) = −61.41 ± 7.13), providing a better fit for ≥ 20/22 subjects in all three environments. When compared against the truncated single-exponential fit, the double exponential model still produced smaller AIC values on average (mean ΔAIC Double Vs Single (fitted to truncated timecourse) = −21.55 ± 5.90), and remained the preferred model for approximately half the subjects. Crucially, the double exponential model was preferred not only for its automated nature (requiring no manual truncation) but also for the magnitude of improvement: when the single model was superior, the advantage was marginal (ΔAIC = 9.7 ± 1.4), whereas the double model’s advantage was substantial (ΔAIC = 52.8 ± 8.9). We therefore used the double exponential fit for further analysis.

Using these fittings we assessed the ‘learning settling time (LST)’—defined here as the time at which participants achieved stable performance in each environment and quantified as the time point at which performance stabilised to ± 10% of the Final Hit Rate (FHR). For the 22 listeners, LST (Figure 2B; Supplemental Figure 2A-C) was similar across all rooms (a total of 180 trials per room and average trial length of 1.5s): Lecture Room: [45.45 ± 41.70 s]; Open-Plan Office: [58.46 ± 32.34 s] and Car Park: [64.52 ± 34.47 s], with a one-way ANOVA revealing no statistical differences in LST for the three reverberant environments [F(2,42)=2.33, p=0.11, *η_p_^2^* = 0.10] suggesting listeners learned/adapted to these environments at the same rate.

We next wondered whether the FHR achieved by a participant within an environment was related to the time point at which LST was reached. To this end, a Pearson’s correlation analysis revealed FHR and the timepoint at which stable performance was achieved were negatively correlated for the Lecture Room: r_(22)_ = −0.66, p < 0.001, 95% CI [-0.86, −0.31] and Open Plan Office r_(22)_ =-0.50, p = 0.02, 95% CI [-0.74, −0.20], but not for the more-highly reverberant Car Park r_(22)_ = −0.08, p = 0.73, 95% CI [-0.44, 0.35], Figure 2C. This analysis suggests that the earlier in time a listener achieves asymptotic performance (i.e., the time point defined as global learning)—presumably by accumulating knowledge about these environments over time—the higher their performance (defined by FHR) in the task, at least in the two less-reverberant environments assessed here.

We analysed learning effects observed in the rANOVA, specifically, whether speech understanding—quantified in terms of d’—improves as the length of the ‘carrier phrase’ was increased (Figure 2D). Consistent with Brandewie and Zahorik (2010; 2013), we found a significant main effect of length of ‘carrier phrase’: [rANOVA: F (3,63) =26.59, p<0.001, *η_p_^2^* = 0.56]. Bonferroni-corrected *post-hoc* pairwise comparisons revealed a significant effect for most carrier phrase lengths (Supplemental Table 1) other than between the two longest, CP2 *vs*. CP3. This plateau in performance between CP2 and CP3 suggests an upper limit in the ability to exploit/accumulate information in reverberant listening environments with increasing sound duration, consistent with Brandewie and Zahorik’s (Brandewie & Zahorik, 2013) observation of a plateau (and a decline in some reverberant environments) in listening performance with increasing length of carrier phrase, and with the existence of an upper limit to exploiting prior exposure to sound environments to benefit listening (Hicks & McDermott, 2024; McDermott et al., 2013; McWalter & McDermott, 2018).

We hypothesized that if prior exposure to the statistics of reverberant rooms arises from an increase in the length of the carrier phrase, then performance to |Number| will always be better than to |Color| for the shortest carrier phrase (CP0: ‘|Color| |Number| now’; Figure 2E) as |Number| is subject to a longer preceding phrase than |Color|. Assessed in terms of reporting |Color| and |Number| alone, listeners performed significantly better (i.e., showed greater sensitivity) for |Number| compared to |Color|: main effect of target word: [rANOVA F (1,21) =48.79, p<0.001, *η_p_^2^* = 0.70], [mean difference = 0.38, p < 0.001] despite there being twice as many (eight compared to four) possibilities. Moreover, a significant interaction ‘carrier phrase length’ x ‘target word’ was observed: [rANOVA F (1,21) =4.33, p=0.008, *η_p_^2^* = 0.17]. *Post-hoc* pairwise comparisons (see Supplemental Table 2) indicate that for all carrier phrases except CP3, performance to |Number| was significantly higher than for |Color|. This result also suggests that a roll-over effect i.e., a lack or limit to the improvement in performance is evident for carrier phrases longer than CP2 across environments (Brandewie & Zahorik, 2013; McWalter & McDermott, 2018).

A potential explanation for poorer performance in reporting |Color| relative to |Number| for the shortest carrier phrase (CP0) is the impact of the utterance of |Color|, namely its immediate appearance at the start of the phrase i.e., the impact of “order/certainty or predictability” in statistical learning (Conway et al., 2010; Daikoku & Yumoto, 2023). One way to determine whether |Color| presented in the context of the CP0 is at all subject to statistical learning (and is therefore not solely reliant on the short-term accumulation of information within a CP0 trial; see Figure 3G) is to assess whether performance for |Color| for the shortest carrier phrase (CP0) improves from the longer-term accumulation of knowledge i.e., the process of meta-adaptation (Robinson et al., 2016) in which adaptation to short-term statistics improves with repeated exposure to those statistics. Specifically, we tested the hypothesis that, independent of the environment, performance for |Color| in later *steady* trials (here, trials 9-10) of the shortest carrier phrase, CP0, are better compared to *initial* (1-2) trials (Figure 2F). Across the total of 360 trials (across carrier phrase lengths and environments), CP0 initial trials occurred early (trial 6±4), while steady CP0 trials (9th–10th occurrences) did not emerge until later (trial 38±7). If this hypothesis is supported, it suggests performance for |Color| for CP0 benefits from meta-adaptive information conveyed in later trials as knowledge about the global structure of the environment accumulates over time.

**Figure 3.**
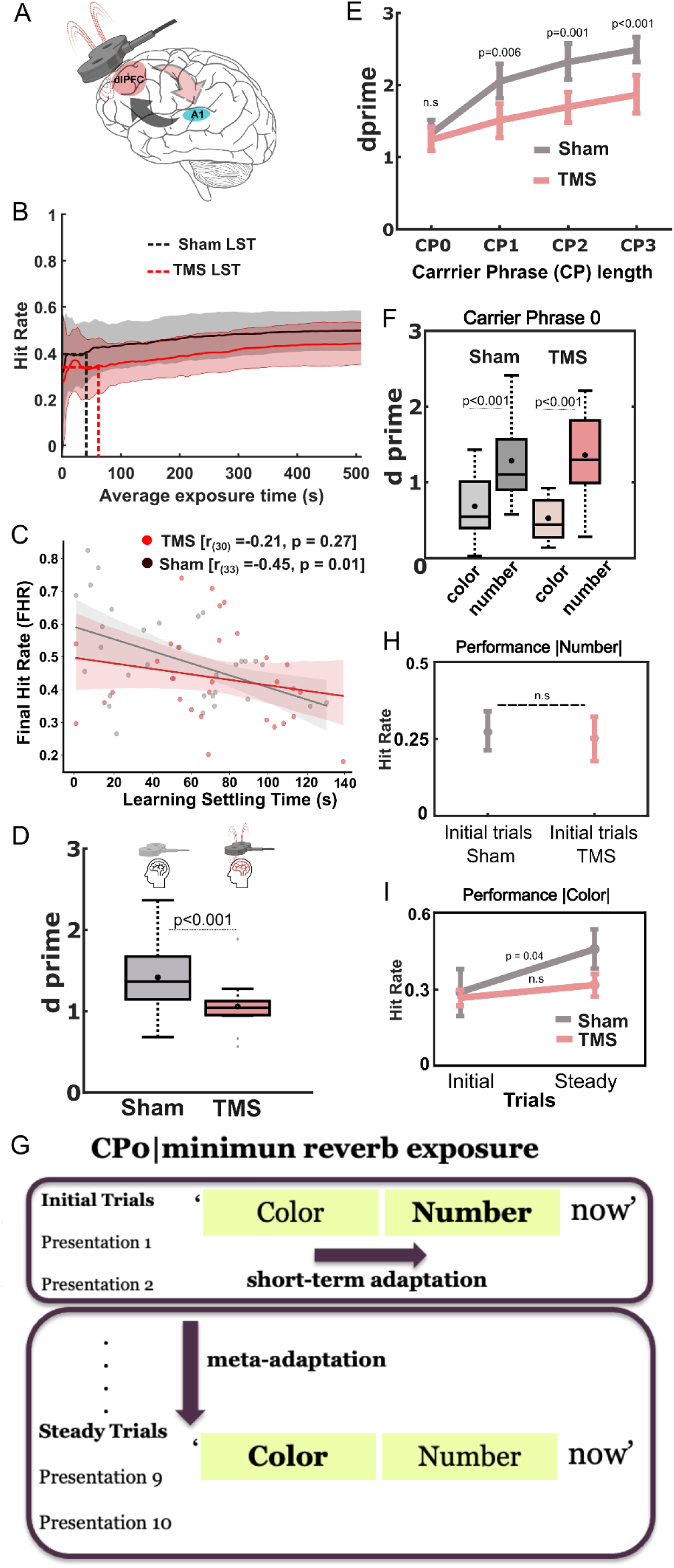
Learning effects in ‘sham’ and ‘TMS’ conditions. **A.** Schematic representation of functional connections between the dorsolateral prefrontal cortex (dlPFC) and primary auditory cortex (A1) under transcranial magnetic stimulation (TMS). Although the TMS protocol was intended to target the dlPFC, it likely affected adjacent prefrontal regions, leading to diffuse downstream effects that may have included modulation of A1**. B.** Average exposure time (s) ‘sham TMS’ and ‘real TMS’ conditions. Correct responses across time spent in each room for ‘sham’ and ‘TMS’ conditions. Solid curves in pink and gray correspond to mean data from ‘TMS’ and ‘Sham’ conditions respectively (after 5-time point moving averages, equivalent to ∼7s, were applied), with the associated shaded areas representing standard errors of the mean. Dashed black (sham TMS) and pink (real TMS) lines represent the time point at which participants reached a stable performance in each environment [i.e., ± 10% of the Final Hit Rate (FHR)], here ‘global learning’, no statistical differences were found for ‘global learning’ between TMS conditions. **C.** Shows a correlation analysis between FHR and the time where 10% of the FHR is achieved in under TMS (pink) and Sham (gray) conditions. Significant negative correlations (Pearson) were observed under Sham stimulation, suggesting the earlier participants reach a stable performance, the higher their FHR is. This relationship was disrupted (lack of significant correlation) under TMS conditions. **D.** Performance in TMS and Sham exposed participants i.e., collapsed sensitivity (d’) to |Color| and |Number| for all carrier phrases and environments (significant main effect of TMS conditions (between-subject’s factor): mixed ANOVA). d’ values plotted in bright pink corresponds to the ‘real TMS’ condition whereas the ‘sham TMS’ accuracy is plotted in gray. Mean d’ is denoted as circles in the boxplot (n=11 for ‘sham TMS’ conditions and n=10 for ‘real TMS’ conditions). **E.** Significant Interaction ‘TMS condition’ x ‘carrier phrase length’. d’ was significantly better as the ‘carrier phrases length’ increased for ‘sham TMS’ compared to ‘real TMS’ except for CPO (n.s), i.e., our proxy for short-term adaptation and minimal exposure to acoustic environments. **F.** Performance to |Color| and |Number| for CP0 i.e., our proxy for short-term adaptation and the carrier phrase where exposure to an environment remained minimal. Here all the environments have been collapsed as only a main effect of “target word”, performance to number always significantly higher than for colour independent of TMS condition. **G.** Schematic for assessing short-term adaptation (performance to |Number| in the Initial Trials) and meta-adaptation (performance to |Color| between Initial and Steady Trials). **H.** Isolated short-term adaptation effects for performance to |Number| in CP0. Hit Rate for Initial (1-2) trials only for |Number| in the CP0 condition for both TMS (pink) and Sham (gray) conditions. Notice that no differences were observed between performance |Number| under Sham or TMS conditions, suggesting no disruption of short-term adaptation. **I.** Hit Rate for Initial (1-2) and Steady (9-10) trials only for |Color| in the CP0 condition for both TMS (pink) and Sham (gray) conditions. Notice that only a significant improvement in performance for from Initial to Steady trials was observed in the ‘sham TMS’ condition. All data available: https://doi.org/10.25949/24295342

Consistent with this hypothesis, a Wilcoxon signed rank test revealed significant better performance (n=22, Z= −2.16, p=0.03, medium effect size (r=0.46)) for |Color| for *steady* trials (9-10; assumed to reflect a meta-adaptive state; Robinson et al., 2016) compared to *initial* trials (1-2; i.e., short-term adaptation). Interestingly, in anechoic conditions i.e., in the absence of reverberation, this meta-adaptive process—the expected improvement in performance between *initial* and *steady* trials—was not observed: Wilcoxon signed rank test [n=10, Z= −0.71, p=0.48, small effect size (r=0.23)], with *initial* trials also appearing early in the task (6±3) and *steady* trials later (36±8). This suggests that performance for |Color| conveyed in the shortest carrier phrase, CP0, improves over time even in the absence of immediate information in the form of any preceding carrier phrase, with knowledge of the statistical/acoustical properties of environments accumulating over the course of the experimental task.

### Improvements in performance are explained by exposure to the environment not talker idiosyncrasies

Normal-hearing listeners are reported to understand speech slightly better when listening to male talkers in a mixture of male and female talkers (Larsby et al., 2015), and quickly adapt to talkers’ idiosyncrasies such as non-native accented speech (Idemaru & Holt, 2011, 2014; Liu & Holt, 2015). Here, we wondered if exposure to the different talkers (3 female and 3 male) could explain the improvement in speech as the length of the carrier phrase was increased in the 22 listeners (Figure 2D). We performed a rANOVA with rooms, carrier length and talker identity as within-subject’s factor. Despite listening performance to some talkers (Supplemental Figure 3A) appearing better than for others [main effect of talkers: [rANOVA: F (5,105) = 27.19, p<0.001, *η_p_^2^* = 0.56], i.e., significantly worse performance for Talker 1 (male) and Talker 6 (female), overall differences in performance due to a specific talker did not explain the benefit to speech understanding as the length of the carrier phrase was increased in each environment, i.e., lack of significant interaction: or ‘talker’ x ‘carrier’: [F (15,315) = 0.06, p=0.84, *η_p_^2^* = 0.028], (see Supplemental Figure 3 for details). This lack of interaction suggests that the rate of improvement in performance with carrier phrase length is uniform across all six talkers.

To explore further the possibility of any talker-specific effects on performance (Best et al., 2008), we assessed the potential benefit of experiencing the same talker on consecutive trials across an experiment, hypothesizing that, independent of room characteristics or length of carrier phrase, experiencing the same talker on consecutive trials would provide a benefit to listening performance. Across all our participants, a total of 1262, 217, and 40 trials were identified as having two, three, or four consecutive trials in which the same talker appeared (see Supplemental Figure 4). Despite the possibility that sustained experience of the same talker might lead to improved listening performance, a Wilcoxon signed rank test revealed no significant differences in performance between trial 1 *vs*. 2 consecutive/same talker trials (n=1262, Z= −0.42, p=0.68, small effect size (r= 0.01)), trial 1 *vs*. 3 consecutive/same talker trials (n=217, Z= −0.17, p=0.87, small effect size (r= 0.01)) or trial 1 vs. 4 consecutive/same talker trials (n=40, Z= −0.159, p=0.11, small effect size (r= 0.02)). Our data suggest that listeners do not use talker identity, at least in the task reported here, to benefit their speech-in-noise understanding.

We also explored the extent to which a talker and/or the task itself could be learned by analysing improvements in performance with increasing exposure to carrier phrases and talkers in the absence of reverberation i.e., under anechoic conditions. In a follow up experiment we assessed performance [similarly to our 3 initial rooms (Brandewie & Zahorik, 2013)] but swapping the Lecture Room with an Anechoic room. Here, in 10 naïve listeners, we isolated only the Hit Rate in the Anechoic room and performed an rANOVA with only carrier phrase length as the within-subject’s factor. Although a significant rANOVA was observed: F (3,27) = 9.10, p=0.007, *η_p_^2^* = 0.57; post-hoc pairwise comparisons, with Bonferroni corrections, revealed that only a significant improvement in performance was observed between CP0 and CP2 [mean difference = 15.67, p = 0.02; see Supplemental Table 4], suggesting that in anechoic conditions very little improvement in performance is achieved by learning the talker/noise/location/task as length of carrier phrase increases. These data support us employing a listening task with high ecological relevance—comprehending speech—to demonstrate the potential benefits of learning background acoustic features to leverage listening performance without requiring listeners to attend to or report abstract features related to the statistics of presumed background sounds *per se* (e.g., Agus et al., 2014, McWalter & McDermott 2018, Bianco et al., 2020).

### TMS disrupts long-but not short-term adaptation to an environment’s reverberation profile

The capacity for learning the statistical structure of acoustic environments and use this knowledge to better understand speech embedded in background noise, suggests a real-world benefit to this form of learned listening. To determine possible brain mechanisms underlying this ability, we reversibly impaired dorsolateral prefrontal cortex (dlPFC, Figure 3A) —a brain region implicated in listening performance in noise—using repetitive transcranial magnetic stimulation (rTMS) and then assessed the ability of listeners to recall |Color| and |Number| in our modified CRM sentences.

The procedure began with TMS manipulation, and although the behavioural task lasted 45 minutes, the inhibitory effects of TMS extended for at least 60 minutes post-stimulation (Gamboa et al., 2010; Hoogendam et al., 2010; Huang et al., 2005; Romero et al., 2022). Specifically, 10 naïve, normal-hearing listeners were subjected to ‘real’ TMS—continuous theta burst on dlPFC for 40s on each side—and then transferred to the anechoic chamber, where they were presented sequences of CRM phrases of varying phrase length in background noise convolved with one of the three listening environments (Lecture Room, Open-Plan Office, and Underground Car Park) as before. A second, control, group of 11 naïve participants underwent ‘sham’ TMS stimulation in which an otherwise-identical TMS procedure was performed with a ‘sham’ TMS coil. All participants were naïve to differences in the TMS procedure as well as to the listening task; indeed, as with the experimental group, participants in the ‘sham’ group experienced a short period of single-pulse TMS stimulation to obtain motor thresholds for involuntary finger movements, followed by ‘sham’ TMS with the stimulator positioned over dlPFC. This process familiarised all participants with the procedure and the influence of TMS in generating involuntary finger movements through direct stimulation of motor cortex. The ‘sham’ procedure was designed to maintain participant blinding, such that participants in the ‘sham’ group would not know whether the subsequent speech-recall task followed active or ‘sham’ stimulation.

Given the potential placebo effects of a ‘sham’ TMS stimulation, we first tested whether our sample of 11 ‘sham’ TMS participants exhibited similar behavioural performance to the larger sample of 22 participants who had not been exposed to any TMS manipulation. A mixed ANOVA with a between-group factor of Condition and within-group factors of room and CP length confirmed that performance in these two populations was comparable, with no significant main effect of TMS conditions observed (‘sham’ *vs*. no exposure to TMS): [F (1,31) =0.01, p=0.91, *η_p_^2^* = 0.00], and no significant interaction Condition x CP length was observed: [F (3,93) =0.48, p=0.69, *η_p_^2^* = 0.01]; confirming that participants experiencing ‘sham’ TMS did not perform significantly differently from the ‘no exposure to TMS’ population.

We next assessed learning over the time-course of the task itself—‘Learning Settling Time—as the cumulative mean hit rate for listeners reporting |Color| and |Number|, fitted for each TMS condition and environment with a two-phase association model (Figure 3B and Supplemental Figure 5 and 6). Our metric for ‘Learning Settling Time’ was achieved for Lecture room at: ‘sham’ TMS [40.22 ± 34.17 s] and ‘real’ TMS [64.86 ± 17.08 s]; for Open-Plan Office: ‘sham’ [51.50 ± 33.26 s] and ‘real’ TMS [67.32 ± 37.04 s]; and for Car Park: ‘sham’ [64 ± 39.02 s] and ‘real’ TMS [69.80 ± 37.31 s]. A mixed ANOVA between ‘sham’ TMS and ‘real’ TMS participants revealed a between-subject’s effect of TMS Conditions: [F (1,19) = 184.42, p<0.001, *η_p_^2^* = 0.91], suggesting participants always achieved an early LST under ‘sham’ stimulation [51.91 ± 36.11s] compared to longer LST under ‘real’ TMS [71.76 ± 36.09 s]. No significant Rooms or Rooms x TMS condition effects were observed.

Pearson’s correlation analysis (collapsed across all environments) revealed a negative correlation between FHR and ‘Learning Settling Time’ for ‘sham’ TMS listeners (see Figure 3C): [r_(33)_ =-0.45, p = 0.01, 95% CI [-0.70, −0.11] but not for ‘real’ TMS listeners: [r_(30)_ = −0.21, p = 0.27, 95% CI [-0.53, 0.16]. Specifically, participants undergoing ‘sham’ TMS achieved stable performance relatively early in the task and maintained this level of performance throughout. Relative delay in attaining stable performance was associated with a lower FHR, with the time at which early LST is achieved predicting overall task performance. This relationship was not evident in listeners subject to ‘real’ TMS, where the time to reach a stable performance was not predictive of the magnitude of FHR. Under TMS manipulation of dlPFC, LST had no predictive value, indicating that task performance was less stable and less reliable for participants exposed to ‘real’ compared to ‘sham’ TMS.

We next employed a mixed ANOVA to compare performance (d’), (see Figure 3D)—of ‘sham’ participants with those who received ‘real’ TMS, and observed a significant between-subject’s effect of TMS condition: ANOVA [F (1,17) = 6.78, p=0.02, *η_p_^2^* = 0.28], where ‘sham’ participants showed significantly better abilities to recall |Color| and |Number| compared to participants subjected to ‘real’ TMS [*post-hoc* pairwise comparisons with Bonferroni corrections: [mean difference = 0.46]. In addition, a significant interaction ‘TMS condition’ x ‘carrier phrase length’ was observed, Figure 3E: [F (3,51) = 6.72, p<0.001, *η_p_^2^* = 0.28]. A *post-hoc* pairwise comparison with Bonferroni corrections showed that performance in the CP0 condition was not statistically different between ‘sham’ and ‘real’ TMS conditions (mean difference = 0.14, p = 0.49), however performance was better for ‘sham’ compared to ‘real’ TMS for CP1 (mean difference = 0.47, p = 0.04), CP2 (mean difference = 0.51, p = 0.02), and CP3 (mean difference = 0.71, p < 0.001). Listeners exposed to ‘sham’ TMS retained the improvements of performance as the length of carrier phrase was increased whereas those listeners receiving ‘real’ TMS—in which activity in dlFPC is presumably impaired—appeared unable to accumulate over time knowledge of the sound environment.

The different time courses over which statistical learning might occur suggests the involvement of multiple neural mechanisms and brain circuits (Anderson & Malmierca, 2013; Antunes & Malmierca, 2011; Robinson et al., 2016), with different cadences of learning potentially controlled, or at least modulated, by different brain centres. The time-course of midbrain neurons to adapt to different sound environments, for example, is slowed, and the capacity for retaining a ‘memory’ of those sound environments disappears, when cortex is inactivated through cooling (Robinson et al., 2016). This suggests feed-forward and feed-back mechanisms, with their own time-courses or time-constants, contribute to performance, including the rate at which sound environments are learned. We wondered if ‘real’ TMS had a detrimental effect on short-term adaptation i.e., the observed advantage in performance for |Number| compared to |Color|, for the shortest carrier phrase CP0 (Figure 2E and 3F). A mixed ANOVA revealed a main effect of ‘target word’, where performance for |Number| was always significantly better than for |Color| for all the environments explored and across TMS conditions [F (1,17) = 31.99, p<0.001, *η_p_^2^* = 0.65]. However, no interaction between ‘TMS conditions’ and ‘target word’ was observed, indicating that in both ‘real’ and ‘sham’ TMS conditions, the same trend of better performance to |Number| compared to |Color| was observed (Figure 3F).

To dissociate specifically short-term adaptation from longer-term meta-adaptation, we compared performance (Hit Rate) to |Number| for *initial* trials (1-2), for CP0 only, as performance for this length of carrier phrase should be influenced only by short-term accumulation of information (i.e., within trial; see Figure 3G). Note that for both groups, CP0 *initial* trials occurred early in the task (360 total trials): trial 5±4 for ‘sham’ TMS and trial 10±9 for ‘real’ TMS. However, *steady* trials (9th–10th instances) did not emerge until trial 34± 6 and 38± 12, respectively.

A non-parametric Wilcoxon signed rank test revealed no significant differences in |Number| performance for *initial* trials between ‘sham’ and ‘real’ TMS exposed listeners (n=10, Z= −0.28, p=0.78, with a small effect size (r= 0.01); Figure 3H), suggesting that rapid learning of the statistical structure of the reverberant environment—in the order of a few hundreds of milliseconds—is resistant to the effects of ‘real’ TMS applied to dlPFC. To determine how much meta-adaptative improvement in performance could be disrupted by applying ‘real’ TMS to dlPFC, we then compared *initial* trials (1-2) and *steady* trials (9-10) for CP0—our proxy for meta-adaptation (Figure 3G). We tested the hypothesis that disruption of dlPFC would affect late (meta-) but not early adaptation. To this end, we expected no differences between performance in *initial* trials between ‘real’ and ‘sham’ TMS (as *initial* trials are influenced only by short-term adaptation), and a relative lack of improvement in performance in later, *steady* trials (meta-adaptation influenced) in ‘real’ compared to ‘sham’ TMS. Consistent with our hypothesis, a Wilcoxon signed rank test revealed no significant differences in performance for *initial* trials between ‘sham’ and ‘real’ TMS exposed listeners (n=10, Z= −0.21, p=0.83, with a small effect size (r = 0.01), but improved performance for *steady* compared to *initial* trials (n=10, Z=-2.06, p=0.04, large effect size (r=0.65)) for ‘sham’ TMS only (Figure 3I). These data suggest that performance in |Color| for the shortest carrier phrase, CP0, was disrupted due to impaired accumulation of knowledge about the environments encountered. In turn, performance under ‘TMS’ in *steady* trials (presumed to be influenced by meta-adaptation) was reminiscent of performance observed in initial trials where only immediate knowledge could be used to improve performance. This is reminiscent also of adaptation to sound environments reported *in vivo* (Dean et al., 2005, 2008; Robinson et al., 2016; Wen et al., 2009), where interrupting efferent feedback by cortical cooling impairs the capacity of midbrain neurons to learn the statistical structure of sound environments; though neurons adapt to the short-term statistical structure of a sound environment each time they are exposed to it, they fail to demonstrate the acceleration of adaptation and improvement in (neural) discrimination performance as the same environment is re-encountered, adapting to it only as if exposed to that environment for the first time.

### Statistical learning of room acoustics is tuned to universally experienced reverberation times

Reverberation is a common feature of many acoustic environments—natural and built (Traer & McDermott, 2016). Interestingly, Zahorik and Brandewie (2013) reported that improvements in speech understanding when a ‘simulated room’ is repeatedly encountered were best when listeners experienced moderately reverberant environments. Specifically, we hypothesized that the ability to understand speech in noise depends on exposure to specific values of RT_60_ across our experimental paradigm, including those in the range commonly experienced by human listeners in natural and built environments (Traer & McDermott, 2016). To test this hypothesis, we recruited a new population of naïve participants with no previous exposure to our experimental paradigm and assessed their ability to understand speech in noise using the same CRM *corpus* as before, but in different combinations of acoustic environments i.e., room contexts defined by their RT_60_s.

We first tested the ability of a new sample of 10 naïve participants to recall |Color| and |Number| in Open-Plan Office and Underground Car Park as before, but with the Lecture Room (RT_60iso_ = 0.42 s) swapped out for a more Highly Reflectant elevator lobby (Figure 4) with an RT_60iso_ of 1.55 s—i.e., between that of the Open-Plan Office (0.96s) and the Underground Car Park (2.42s). We therefore changed the ‘room-context’ in which Open-Plan Office and Underground Car Park were learned. When contrasting the performance (Final Hit Rate) in an ANOVA considering performance only in the common rooms (Open-Plan Office and Underground Car Park) of these 10 naïve participants *vs*. the 11 subjects randomly selected from the initial 22 participants sample, we found that speech understanding in Open-Plan Office and Underground Car Park was significantly better when the room acoustics of these environments were learned in the context of the Lecture Room than in the context of the Highly Reflectant Room [mean difference = 14.8, F (1,41) = 30.72, p<0.001, *η_p_^2^* = 0.45].

**Figure 4.**
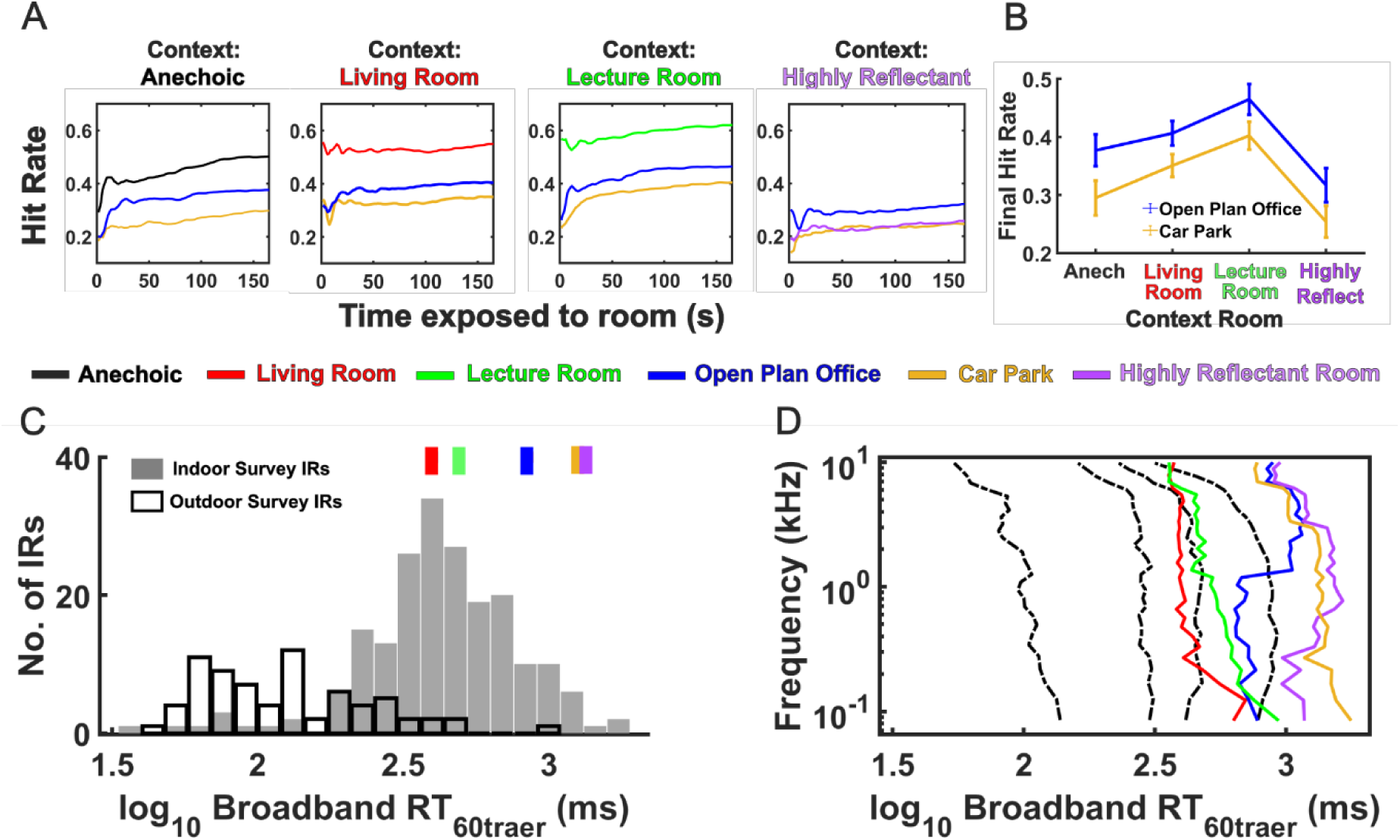
Performance in different 3-room combinations where acoustics of 3 rooms span an ecological range. **A.** Hit rates over time for different 3-room combinations. Mean data for four different 3-room combinations are plotted where Open-Plan Office (*dark blue*) and Underground Car Park (*yellow*) were always included and the context room was either Anechoic (*black*), Living Room (*red*), Lecture Room (*green*), Lecture Room (*green*) or Highly Reflectant (*purple)*. **B.** Performance in Open-Plan Office and Car Park when paired with different context rooms. Final hit rates were highest for Office and Car Park when presented in conjunction with the Lecture Room, followed by the Living Room and Anechoic with poorest performance observed when the Highly Reflectant space was the context. **C.** Environments tested compared to ecological range. The RT_60traer_s for the five test rooms, calculated as the median RT_60_ value across 31 frequency sub-bands (Traer & McDermott, 2016), are plotted (*colored boxes*) above a histogram of the median RT_60traer_s calculated for 199 indoor (*filled bars*) and 72 outdoor (*empty bars*) spaces recorded by Traer and McDermott (2016). **D**. Frequency dependence of reverberation time (RT_60traer_) in test rooms compared to ecological range. RT_60_s of the 5 test rooms are displayed for frequency sub-bands used to calculate RT_60traer_ (*colored lines*). Quartiles for combined indoor and outdoor spaces (Traer & McDermott, 2016) are plotted as *black dashed lines*. Decreasing reverberation time above 0.5kHz is observed for Living Room, Lecture Room and Car Park as has been typically described for indoor spaces (Traer & McDermott, 2016); however RT_60traer_ profiles for Open-Plan Office and Highly Reflectant space were notable for their longer reverberation times at and above 1kHz. All data available: https://doi.org/10.25949/24295342

Wondering whether the ‘room-context’ effect was determined by task difficulty i.e., the more-highly reverberant lobby being intrinsically more difficult for listening than the Lecture Room [mean difference = 35.75, t _(9)_ =18.72, p < 0.001, Cohen’s d = 5.92]. We recruited a further new sample of 11 naïve participants and performed the same task in which they were exposed to another combination of three rooms, here with the Lecture Room swapped for an Anechoic Room. Anechoic spaces—rooms whose walls are treated to completely absorb reflected sound energy—have been extensively reported to aid speech understanding, especially in background noise. To test this hypothesis directly, we recruited a further sample of 10 naïve participants and compared their performance, in our task, in a combination of an Anechoic Room, Open-Plan Office and Underground Car Park, with the performance of 11 subjects randomly selected from the initial 22 participants’ sample. Although task difficulty might be expected to be at its lowest i.e., easiest, in the less-reverberant Anechoic Room, this was not the case, performance in the Anechoic room was poorer compared to Lecture Room [mean difference = 11.08, t _(9)_ =2.66, p = 0.013, Cohen’s d = 0.84] (Figure 4 A-B). Additionally, we found that performance in the common rooms: Open-Plan Office and Underground Car Park, was significantly better when these environments were again learned in the context of the Lecture Room: [mean difference = 9.7, F (1,41) = 13.24, p<0.001, *η_p_^2^* = 0.26].

Although reverberation might be considered detrimental to speech-in-noise performance, short-term benefits of more, compared to less, room reverberation for speech understanding have, in fact, been reported, including for the task we report here (Brandewie & Zahorik, 2010, 2013). Further, it is likely that listening performance in noise is indeed better when listeners have access to more commonly experienced, or ethologically realistic, levels of reverberation. We tested this hypothesis by recruiting a new sample of naïve listeners and swapping again the Lecture room with a Living Room with a shorter RT_60iso_ (0.33 s). When comparing performance between Lecture and Living Room no significant differences were observed: [mean difference = 7.04, t _(10)_ =1.69, p = 0.061, Cohen’s d = 0.51] (Figure 4 A-B). However, significantly better performance was observed in the common rooms: Open-Plan Office and Underground Car Park when these environments are learned in the context of the Lecture room: [mean difference = 9.7, F (1,43) = 5.79, p=0.21, *η_p_^2^* = 0.13]. These results suggest that when certain reverberant characteristics are encountered e.g., Lecture Room, they might unlock a perceptual advantage in understanding speech in harder/more reverberant environments such as Open-Plan Office and Indoor Car Park.

Seeking to explain why some rooms might be better than others in aiding speech understanding, we recalculated reverberation times for our 5 reverberant rooms according to the method of Traer and McDermott (2016) where RT_60traer_ is equivalent to the median RT_60_ measured in 31 sub-bands with centre frequencies between 80 Hz and 10 kHz (see Methods). We then compared these RT_60traer_ values with the reverberation times recorded for a wide range (n= 271) of built and natural environments by the same authors (Figure 4C-D. RT_60traer_s for 5 rooms conditions and Traer’s data, plotted as log_10_ RT_60traer_ for visualisation). In particular, the RT_60traer_ of the Lecture Room, at 0.49 s, lies extremely close to the median and mean RT_60traer_ values of the built environments [0.42 s and 0.50 s respectively generated by the skewed distribution (n= 199), Figure 4C]. This suggests that superior performance in understanding speech understanding in noise in the Lecture Room—and the positive impact on speech understanding when the Lecture Room is included in the three-room listening task—is related to its reverberant characteristics being commonly encountered in everyday listening situations. Notably, the mean and median RT_60traer_s of the recorded natural environments were much lower (0.12 s and 0.16 s, respectively [n = 72], Figure 4C) than the RT_60traer_s of the Lecture Room. Together, these data suggest that brain mechanisms contributing to effective speech understanding in reverberant environments might have adapted to the range of environments experienced over some longer time course than we assessed here.

## Discussion

We assessed the ability of listeners to understand speech embedded in background noise and convolved with the room impulse responses (RIRs) of different indoor environments defined by their reverberation decay time, or RT_60_—the time taken for reverberant sound energy to decay 60 decibels (dB). We found speech understanding improved with repeated exposure to an environment, a form of learning that was impaired following continuous, bilateral theta-burst transcranial magnetic stimulation (TMS) of dorsolateral prefrontal cortex (dlPFC). Specifically, we observed rapid learning i.e., an improvement in speech understanding over the first few seconds of room exposure, likely due to listeners learning the acoustic reverberation—the only variable held constant across trials for a given environment. This learning, on the timescale of several seconds, was impaired by TMS, whilst learning at shorter or longer timescales was not, suggesting that TMS applied to dlPFC specifically disrupted learning of the reverberant characteristics of an environment. Listeners also showed better listening performance in moderately reverberant environments, and this performance was transferable between different acoustic environments. Specifically, the ability to report correctly keywords spoken in environments with the more extreme—higher—RIRs was best when these environments were encountered in experimental blocks also containing the moderately reverberant environments. This transference suggests an ethological tuning to more commonly encountered environments when learning acoustic backgrounds.

### A role for dlPFC in statistical learning of room acoustics

Inactivation of dlPFC using theta-burst TMS reduced performance in our speech-in-noise task, assessed in terms of the hit rate for |Color| and |Number| in CRM phrases as a function of duration of exposure to a sound environment. Notably, however, TMS did not impair listening performance for the shortest-duration phrase, ‘|Color| |Number| now’. For this duration of carrier phrase, performance i.e., the tendency to report more accurately |Number| compared to |Color|—despite |Number| having twice as many options as |Color|—was unaffected by TMS stimulation of dlPFC. The robustness of listening performance to the shortest carrier phrase suggests that the ability to hear out speech in background noise and potentially to learn this over time in complex acoustic environments does not rely solely on cortical feedback from dlPFC. The ability to leverage performance on |Number| by exposure to |Color| presented in background noise convolved with the room impulse response is evident even when the function of dlPFC is (presumably) impaired. It is the case, however, that the rate of learning—i.e., how rapidly performance improves with increasing time of exposure to an acoustic environment—was influenced when dlPFC was impaired. Specifically, under the influence of prior TMS stimulation, listeners were less able, and less reliable, when learning the acoustic background features of an environment to enhance their speech understanding, particularly in the more challenging environments of the Underground Car Park and Open-Plan Office.

Dense connections between dlPFC and the sensory cortices provide the conduit for prediction-based, top-down regulation of learning (Morrone, 2010). These model-based predictions rely on learned templates of sensory environments (Alexander & Brown, 2018) that are critical for making inferences under situations of sensory uncertainty, such as those encountered in noisy and challenging listening environments (Bartolo & Averbeck, 2021). Dorsolateral prefrontal cortex can influence auditory processing by means of direct connections to auditory cortex (Hackett et al., 1999; Plakke & Romanski, 2014)—see Figure 3Ai—and to the auditory thalamic reticular nucleus, a key modulator of auditory thalamo-cortical loops (Zikopoulos & Barbas, 2006). In return, dlPFC receives projections from primary and secondary auditory cortex (Barbas & Pandya, 1987; Barbas & Pandya, 1991; Goldman-Rakic & Schwartz, 1982; Pandya & Barnes, 2019; Petrides & Pandya, 2002), indicating that a reciprocal auditory-to-prefrontal ‘listening loop’ exists to support hierarchical predictions and prediction-error feedback. These ‘listening loops’ could potentially extend to cortical efferent feedback (Blackwell et al., 2020; McAlpine & de Hoz, 2023) and modulate the function of inner-ear sensitivity in attended speech-in-noise tasks (de Boer & Thornton, 2008; Garinis et al., 2011; Giraud et al., 1997; Mishra & Lutman, 2014; Hernández-Pérez et al., 2021).

The specific mechanism by which TMS-induced modulation of dlPFC impairs listening performance remains unknown. Neurostimulation studies targeting dlPFC during learning paradigms have generated conflicting directions of effect i.e., stimulation of dlPFC is reported to impair implicit learning (Nydam et al., 2018; Pascual-Leone et al., 1996) or enhance it (Ambrus et al., 2020). Ambrus and colleagues argued that impairment of dlPFC during implicit/statistical learning allows for increased engagement of a learning mechanism that is ‘model-free’ i.e., does not rely on inherent sensory representations and therefore takes longer to consolidate. In the present study, although the TMS protocol was designed to target dlPFC using the 10–20 EEG cap system, this approach does not permit precise stimulation of dlPFC (Jiang et al., 2025; Nikolin et al., 2019; Trapp et al., 2020) or its direct projections to the auditory cortex. Instead, stimulation likely influenced prefrontal regions producing diffuse downstream effects including the auditory cortex and other brain regions such as hippocampus (Bennett et al., 2011; Stillman et al., 2013), striatum (Albouy et al., 2015) and basal ganglia (Packard & Knowlton, 2002). We therefore speculate that the short periods over which learning is required in our task are subserved by a pre-frontally mediated, ‘model-based’ learning able to construct predictions ‘on the fly’ (Daw et al., 2005). We suggest, in our study, that disrupting a rapidly acting model-based form of learning supported by prefrontal regions, as part of the auditory-prefrontal ‘listening loop’, may lead to impaired learning of reverberant environments.

Dorsolateral prefrontal cortex is also implicated in several, high-level cognitive functions including multi-sensory integration (Fuster et al., 2000), executive functions such as inhibition and working memory (Castro-Meneses et al., 2016; Coltheart et al., 2018; Wang et al., 2015) and listening in noisy environments (Du et al., 2016). The impact of TMS on performance of our speech-in-noise task may be linked to the involvement of dlPFC in more generalized cognitive functions such as executive, memory, and attention processes. Dissociating the potential influence of TMS on such processes relative to a specific role in speech-in-noise understanding in noisy, reverberant environments would be an ideal next step. Specifically frontal regions have been linked to sensorimotor integration during speech-in-noise perception in young and older human participants (Du et al., 2016), suggesting that TMS might disrupt global functioning required for the analysis and understanding of speech.

Whilst not entirely unrelated, all participants completed a familiarization task preceding the main experiment, which consisted of 10 trials presented in anechoic conditions at 0 dB Signal-to-Noise Ratio (SNR). Participants exposed to either ‘real’ or ‘sham’ TMS underwent this task following cTBS procedures. Notably, all participants, including those subjected to cTBS, achieved 75-100% accuracy during the familiarization phase, with no statistical differences between TMS groups [mean difference=1.25, t_(1, 9)_=0.43, p=0.68, d=0.14], confirming their comprehension of the task and successful procedural learning (i.e., learning the task *per se*). In addition, speech understanding in the most challenging condition (CP0) was not disrupted following TMS manipulations, suggesting that the effect of TMS was specifically in terms of influencing the way these environments were learned over time. Nevertheless, we acknowledge that the primary task may have imposed greater cognitive demands, potentially requiring a more significant involvement of the dlPFC to meet task requirements.

Further, we cannot rule out individual variability in the duration of TMS effects, nor differences in effect duration across anatomical sites—such as the motor cortex (Gamboa et al., 2010; Hoogendam et al., 2010; Huang et al., 2005; Romero et al., 2022) compared to the dLPFC examined in our study. Continuous theta burst stimulation (cTBS), as used in this study, typically induces aftereffects lasting 20–50 minutes (Huang et al., 2005; Wischnewski & Schutter, 2015). While these effects are well established in the motor cortex, where motor-evoked potential changes can persist for up to one-hour, comparable durations of cortical modulation have also been reported in the prefrontal cortex (Taylor et al., 2024; Wagner et al., 2006). Specifically, inhibitory rTMS applied to the dorsolateral prefrontal cortex (dlPFC) has been shown to reduce cortical oxygenation levels—indicative of decreased neural activity—for at least 45 minutes (Tupak et al., 2013). Given that changes in cerebral haemoglobin concentration closely correspond to neuronal activation (Liao et al., 2013), these findings support the use of functional Near Infrared Spectroscopy (fNIRS) as an indirect but reliable method to infer TMS-induced neural modulation within this timescale. Although no objective post-stimulation measures were collected in the present study, we acknowledge this as a limitation and plan to incorporate such measures in future work to confirm the duration and magnitude of TMS effects more directly.

### What is being learned in statistical learning of acoustic features?

The need to communicate in reverberant environments is common to daily life. It is well known that long reverberation times have detrimental effects on speech quality and intelligibility (Brandewie & Zahorik, 2010, 2013; Srinivasan & Zahorik, 2012; Pavel Zahorik & Brandewie, 2016) and that, specifically, reverberation blurs phoneme boundaries, increasing the extent to which similar words might be confused, e.g., ‘sir’/’stir’ (Watkins, 2005b; Watkins & Makin, 2007). However, with sufficient exposure to longer reverberation times in the form of a carrier phrase, similar levels of word identification to those achieved in minimal reverberation are observed (Watkins, 2005b; Watkins & Makin, 2007). Based on the premise that providing sufficient contextual information enables the auditory system to adapt and compensate for the detrimental effects of reverberation, Zahorik and colleagues (Brandewie & Zahorik, 2010, 2013; Srinivasan & Zahorik, 2012; Zahorik & Brandewie, 2016) and, here, ourselves, demonstrate that prior exposure to the reverberant characteristics of a room can enhance sentence understanding when that environment is re-encountered. Moreover, Zahorik and colleagues, and our own data (see Figure 4A), demonstrate that improvements in speech understanding are always greater in reverberant compared to anechoic, environments (Brandewie & Zahorik, 2010, 2013; Srinivasan & Zahorik, 2012), i.e., improved performance arises only through exposure to a reverberant environment rather than to the speech material *per se*.

Adaptation to continuous noise is a well-known phenomenon in the auditory system (Costalupes et al., 1984; Gibson et al., 1985; Phillips, 1985; Rees & Palmer, 1988) that could potentially also explain the speech improvements observed among carrier lengths. This is particularly relevant in our experiments because the masking noise preceded the onset of the speech material, potentially providing a window for noise-adaptation to occur (Ainsworth & Meyer, 1994; Ben-David et al., 2016). However, Zahorik and Brandewie reported similar effect sizes in speech enhancements when 1 s (Brandewie & Zahorik, 2010) or 150 ms (Brandewie & Zahorik, 2013) of white noise was presented prior to the carrier phrase (i.e., CP0—|Color| |Number|). From this, the authors concluded that noise alone did not convey sufficient information about the acoustic environment to elicit improvements in speech understanding.

Adult listeners can also acclimatise to speech acoustics that deviate from the norm or from long-term language regularities such as dialects and foreign accents, with sufficient exposure (Idemaru & Holt, 2011, 2014; Liu & Holt, 2015). Even if our participants were all Australian-English native speakers exposed to an American-English speech corpus—the CRM—some perceptual learning of individual talker’s idiosyncratic speech patterns might arise over the course of the task (Choi & Perrachione, 2019; Liu & Holt, 2015; Stilp, 2020). In addition, it has been reported that ‘target voice continuity’ i.e., the same talker within trials—similar to our design—can enhance the build-up of selective attention and improve listeners abilities to report correctly digits sequences in the presence of other talkers (Best et al., 2008). Because talker identity did not vary randomly within individual sentences in our paradigm, we cannot completely rule out that some degree of within-trial voice tuning occurred equally across the six talkers during the carrier phrase. However, three lines of evidence indicate that voice tuning alone cannot account for our findings. First, when the room’s build-up effect—the effect assessed here—is explored in an anechoic space using the exact same talkers, there is markedly reduced improvement in speech understanding with increasing the length of the carrier phrase (Brandewie & Zahorik, 2010, 2013, Supplemental Table 4) and no longer-term meta-adaptation across trials (Figure 2G). Second, speech understanding for the exact same voices in the Open-Plan Office and Car Park varied significantly depending solely on the reverberation time of the third context room in the block. Third, interrupting the continuity of the room—its reverberation profile—rather than the continuity of the target talker, is the factor reported to disrupt significantly the improvement in speech understanding with increasing exposure (Brandewie & Zahorik, 2018). Thus, while rapid adaptation to a talker’s voice may provide a benefit (Cooke et al., 2022; Kakehi, 1992; Kato & Kakehi, 1988) the substantial gains in performance from increasing length of the carrier phrase, meta-adaptation, and room-context effects observed here are driven primarily by learning the reverberant environments.

A key feature of this study, one that distinguishes it from previous assessments of statistical learning of acoustic features in human listeners, is our use of an ethologically valid listening task—understanding speech in background noise while listeners implicitly learned background features. Given the perceptual biases inherent in sensory processing, it is notable that investigators have previously reported different noise tokens to be differently learnable (Agus et al., 2014; Daikhin et al., 2017), for example, and that textures must be carefully controlled to ensure listeners are not exploiting subtle spectro-temporal features in their judgments of statistical similarity (McDermott et al., 2013). The potential for listeners to hear out specific—potentially unique to them—spectro-temporal features of otherwise statistically identical sound tokens, or to make judgments on the regularity or similarity of tone sequences based on unique listening experiences (Barascud et al., 2016; Bianco et al., 2020), presents a potential confound for these studies. Our study countermands this problem by actively exploiting the propensity for human listeners to ascribe meaning to spectro-temporal fluctuations (here, speech) in acoustic waveforms (Brandewie & Zahorik, 2010, 2013). Though still targeting the learning of background acoustic features, it engages listeners in a highly relevant listening task, one for which humans have rapidly evolved—i.e., understanding speech in background noise. Listeners’ attention, therefore, was called away from the background acoustic features to the ethologically relevant foreground task of attending to human speech, void of semantics. Additionally, length of phrase, talker, and |Callsign| were uninformative to |Color| and |Number| in our utterances, as were |Color| and |Number| to each other. This misdirection allows us to exploit speech as a ‘reporter’ or biomarker for statistical learning of background, more-abstract, acoustic features, and without the potential confounding factor of (individualised) perceptual bias. Further, we have demonstrated that prior exposure to ethologically relevant environments matters when learning less-commonly encountered acoustic scenes.

The impact on speech understanding of more extreme (longer RT_60_s) environments is reminiscent of the detrimental effect of TMS, with lower listening performance. This suggests that learning the statistical structure of sound environments, to aid speech understanding in reverberant background noise, requires some longer form of memory (experience, developmental, or evolutionary). Brain circuits, therefore, might act to improve listening performance, operating best when at least some of the exposure to listening environments includes plausible, and commonly experienced, reverberation times (Traer & McDermott, 2016). When the acoustics of commonly encountered rooms were replaced with those of other environments having higher or lower amounts of reverberation, performance declined. Intriguingly, the most effective reverberation time we employed, at least in terms of its capacity to be learned and potentially from which learning could be transferred—the RT_60traer_ of 0.49 s of the small Lecture Room is very close to the median and mean RT_60traer_ of 0.42 and 0.50 s previously recorded for a wide range of built environments (indoors and outdoors). This suggests that statistical learning of acoustic background features might be tuned to ethologically relevant environments. Alternatively, built environments might be constructed to generate reverberation subjectively most suited to maximising speech-in-noise understanding, though we are unaware as to whether such a constraint is consciously applied in the design of listening spaces given contingencies such as the range and relative consistency and constancy of potential contents of any given space that influence its reverberant characteristics. Compared to built environments, however, the RT_60traer_s of natural environments were much lower than all the rooms we assessed, save for that of the anechoic room, which was suboptimal in terms of learned listening performance in our task.

### Neural adaptation as a contributing mechanism to the learning of room acoustics

Adaptation to stimulus statistics has been proposed as a neurophysiological mechanism underlying selective suppression of background noise in auditory cortex (Fuglsang et al., 2017; Kell & McDermott, 2019; Khalighinejad et al., 2019; Mesgarani et al., 2014). More recently, it has been shown that spectro-temporal receptive fields of auditory-cortical neurons are sensitive to the RT_60_s of reverberant noise i.e., a form of adaptation specific to reverberation and consistent with a de-reverberation of the environment encountered (Ivanov et al., 2022). Cortical adaptation to reverberation has been also reported in awake listeners (Fuglsang et al., 2017; Mesgarani et al., 2014) and to this end, our data suggest a role for top-down mechanisms in improving over time an attended task—recalling spoken words—in reverberant environments. It is therefore possible that goal-directed behaviours and the feedback the auditory cortex receives from areas such as dlPFC may contribute to fine-tuning the adaptation to reverberant environments.

Our data are also consistent with reports of midbrain auditory neurons recorded *in vivo* adapting to the statistical structure of evolving sound environments over the course of several hundred milliseconds (Dean et al., 2005, 2008) and this capacity for adaptation increases with repeated exposures to the same statistically structured environment. Importantly, whilst rapid adaptation is retained when descending cortical influences are disrupted (through cortical cooling), the longer-term learning effect—a speeding up of adaptation referred to as meta-adaptation—is abolished (Robinson et al., 2016). As with auditory neurons recorded *in vivo* (Bakay et al., 2018; Dean et al., 2005, 2008; Ivanov et al., 2022), it seems listeners might adapt to the statistical structure of a sound environment within a few hundred milliseconds of exposure to enhance performance in a listening task, an initial, rapid, learning phase impervious to inactivation of the dorsolateral prefrontal cortex by theta-burst TMS. The nested set of temporal sensitivities to room acoustics we observe is consistent with features of statistical learning that emerges from the level of the auditory nerve to primary cortex *in vivo* (Dean et al., 2005, 2008; Watkins & Barbour, 2008; Wen et al., 2009), and likely modulated by efferent influences that span the entire auditory pathway all the way to the sensory receptors of the inner ear (Hernández-Pérez et al., 2021; Perrot et al., 2006; Terreros & Delano, 2015) and even the mechanical sensitivity to sound of the eardrum and ossicles (middle-ear bones) (Gruters et al., 2018).

Dissociating auditory cortical feedback circuits to the midbrain impairs the capacity of midbrain neurons to ‘recall’ previous experience of that environment (Bajo et al., 2019; Robinson et al., 2016). Like the performance in our listening tasks, neurons were still able to adapt rapidly (within hundreds of milliseconds) each time an environment was encountered as if for the first time. However, they showed no capacity to exploit a memory of that environment to improve neural coding (i.e., meta-adaptation) (Robinson et al., 2016). Meta-adaptation was consistent with neurons in lower brain centres adapting to the current sound environment, and those in higher brain centres learning the longer-term statistical structure of changing environments. This suggests that once higher brain centres learn an environment, this information, conveyed to early brain centres, ensures they are ‘primed’ for coding an environment when it re-occurs.

## Inclusion and Ethics statement

This study was approved by the Human Research Ethics Committee of Macquarie University (ref: 5201833344874). Each participant signed a written informed consent form and was given a small financial remuneration for their time.

## Acknowledgements

The study was supported by the Australian Research Council (DP180102524 awarded to D.M, J.J.M.M, and P.F.S, and FL 160100108 awarded to D.M.). The authors would like to thank Jörg Bulcholz and Javier Badajoz-Davila for their assistance with the spatialization and sound field simulation of the acoustic stimuli. We thank Adam Weisser for his assistance in obtaining the room acoustic calculations for the actual rooms used in this study. We would also like to thank Kurt Shulver for his assistance during rTMS manipulations. We sincerely thank Yuranny Cabral-Calderin for her valuable insights and constructive feedback on improving this manuscript.

## Authors’ contributions (CReditT taxonomy: (McNutt et al., 2018))

Conceptualization: H.H-P., J.J.M.M., J.M-H., J.T., P.F.S. and D.M.

Data Curation: H.H-P., J.J.M.M., J.M-H., and D.M.

Formal Analysis: H.H-P., J.M-H. and J.T.

Funding Acquisition: H.H-P., J.J.M.M., P.F.S. and D.M.

Investigation: H.H-P and J.T.

Methodology: H.H-P., J.J.M.M., J.M-H., J.T., P.F.S. and D.M.

Project Administration: H.H-P. and P.F.S

Resources: H.H-P., J.J.M.M., J.M-H., J.T. and P.F.S.

Software: H.H-P., J.J.M.M., J.M-H. and J.T.

Supervision: H.H-P., J.J.M.M., P.F.S. and D.M.

Validation: H.H-P., J.J.M.M., J.M-H., J.T., and D.M.

Visualization, H.H-P., J.M-H.,J.T. and D.M

Writing – Original Draft Preparation: H.H-P., J.M-H., J.T., P.F.S. and D.M.

Writing – Review & Editing: H.H-P., J.J.M.M., J.M-H., J.T., P.F.S and D.M.

## Declaration of interests

The authors declare no competing interests

---

**Supplemental Table 1.** Pairwise comparisons between performance (d’) for carrier phrases length n 22 participants.

| **(I) Carrier Phrase** | **(J) Carrier Phrase** | **Mean Difference (I-J)** | **Std. Error** | **Sig.<sup>b</sup>** | **95% Confidence Interval for Difference<sup>b</sup>** |  |
|---|---|---|---|---|---|---|
|  |  |  |  |  | **Lower Bound** | **Upper Bound** |
| CP0 | CP1 | -.295<sup>*</sup> | .070 | .002 | -.498 | -.091 |
|  | CP2 | -.510<sup>*</sup> | .063 | <.001 | -.692 | -.328 |
|  | CP3 | -.586<sup>*</sup> | .089 | <.001 | -.846 | -.325 |
| CP1 | CP2 | -.215<sup>*</sup> | .045 | <.001 | -.345 | -.085 |
|  | CP3 | -.291<sup>*</sup> | .083 | .012 | -.532 | -.050 |
| CP2 | CP3 | -.076 | .074 | 1.000 | -.291 | .139 |

Based on estimated marginal means
\*. The mean difference is significant at the .05 level.
b. Adjustment for multiple comparisons: Bonferroni.
data available: https://doi.org/10.25949/24295342

**Supplemental Table 2.**
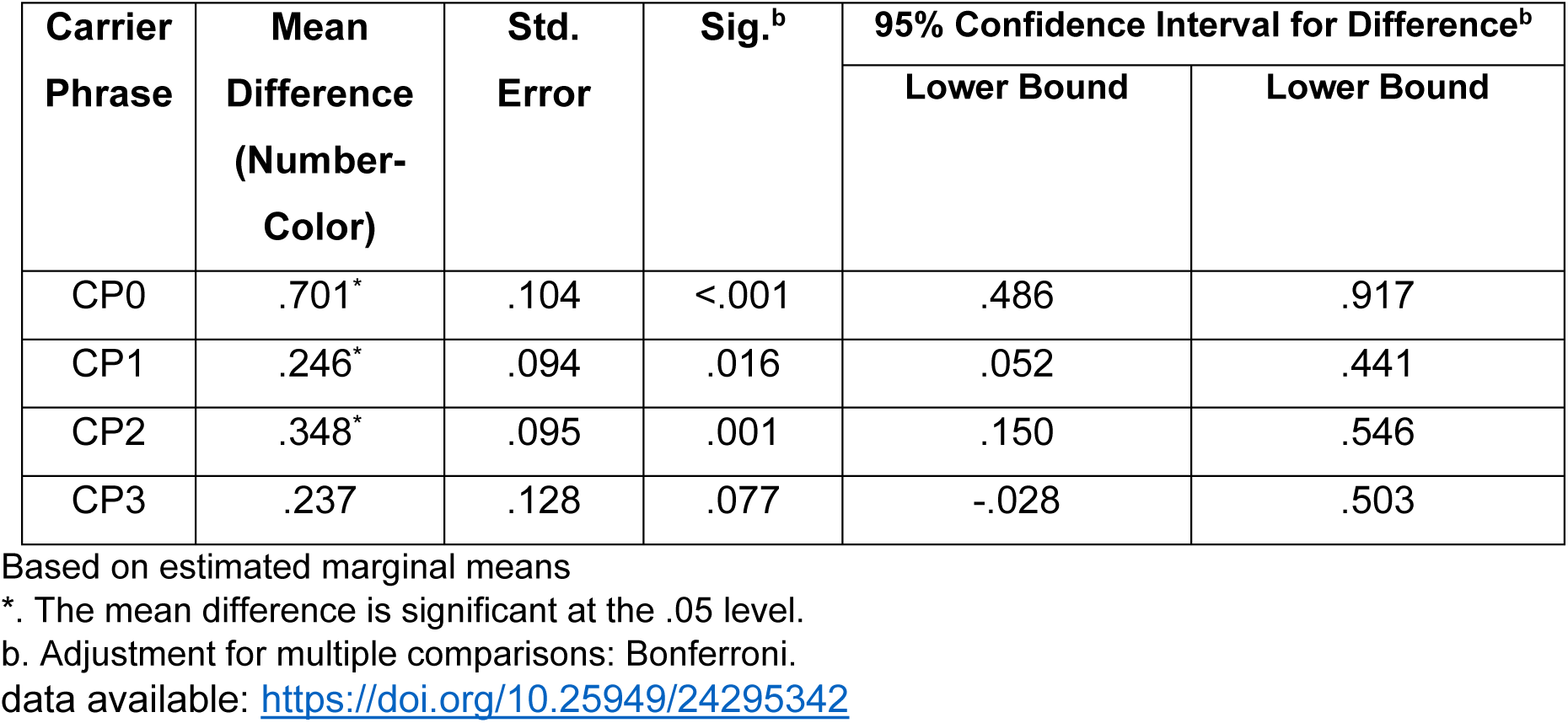
Pairwise comparisons among performance (d’) for Interaction ‘target word’ x ‘carrier phrase length’ in 22 participants.

| **Carrier Phrase** | **Mean Difference (Number-Color)** | **Std. Error** | **Sig.<sup>b</sup>** | **95% Confidence Interval for Difference<sup>b</sup>** |  |
|---|---|---|---|---|---|
|  |  |  |  | **Lower Bound** | **Lower Bound** |
| CP0 | .701<sup>*</sup> | .104 | <.001 | .486 | .917 |
| CP1 | .246<sup>*</sup> | .094 | .016 | .052 | .441 |
| CP2 | .348<sup>*</sup> | .095 | .001 | .150 | .546 |
| CP3 | .237 | .128 | .077 | -.028 | .503 |

Based on estimated marginal means
\*. The mean difference is significant at the .05 level.
b. Adjustment for multiple comparisons: Bonferroni.
data available: https://doi.org/10.25949/24295342

**Supplemental Table 3.** Performance (Final Hit Rate) pairwise comparisons for the six talkers in 22 participants.

| **Talkers** |  | **Mean Difference** | **Std. Error** | **Sig.<sup>b</sup>** | **95% Confidence Interval for Difference<sup>b</sup>** |  |
|---|---|---|---|---|---|---|
|  |  |  |  |  | **Lower Bound** | **Upper Bound** |
| **1** | **2** | -.042 | .016 | .277 | -.096 | .012 |
|  | **3** | -.060* | .013 | .002 | -.101 | -.018 |
|  | **4** | -.104* | .015 | .000 | -.154 | -.055 |
|  | **5** | -.129* | .017 | .000 | -.185 | -.073 |
|  | **6** | .015 | .014 | 1.000 | -.032 | .063 |
| **2** | **3** | -.018 | .014 | 1.000 | -.066 | .030 |
|  | **4** | -.063* | .015 | .007 | -.113 | -.012 |
|  | **5** | -.087* | .015 | .000 | -.137 | -.037 |
|  | **6** | .057 | .017 | .054 | -.001 | .115 |
| **3** | **4** | -.045 | .015 | .123 | -.096 | .006 |
|  | **5** | -.069* | .014 | .001 | -.116 | -.022 |
|  | **6** | .075* | .013 | .000 | .032 | .118 |
| **4** | **5** | -.025 | .017 | 1.000 | -.080 | .031 |
|  | **6** | .120* | .018 | .000 | .060 | .179 |
| **5** | **6** | -.144* | .015 | .000 | -.193 | -.095 |

Based on estimated marginal means
\*. The mean difference is significant at the .05 level.
b. Adjustment for multiple comparisons: Bonferroni.
data available: https://doi.org/10.25949/24295342

**Supplemental Table 4.** Pairwise comparisons between performance (Final Hit Rate) for carrier phrases length in 10 participants under anechoic conditions.

| **(I) Carrier Phrase** | **(J) Carrier Phrase** | **Mean Difference (I-J)** | **Std. Error** | **Sig.<sup>b</sup>** | **95% Confidence Interval for Difference<sup>b</sup>** |  |
|---|---|---|---|---|---|---|
|  |  |  |  |  | **Lower Bound** | **Upper Bound** |
| CP0 | CP1 | -6.000 | 3.602 | .781 | -18.118 | 6.118 |
|  | CP2 | -15.667* | 3.938 | .019 | -28.914 | -2.419 |
|  | CP3 | -13.167 | 4.314 | .082 | -27.679 | 1.346 |
| CP1 | CP2 | -9.667 | 2.885 | .051 | -19.371 | .038 |
|  | CP3 | -7.167 | 2.972 | .235 | -17.164 | 2.830 |
| CP2 | CP3 | -2.500 | 1.537 | .829 | -7.669 | 2.669 |

Based on estimated marginal means
\*. The mean difference is significant at the .05 level.
b. Adjustment for multiple comparisons: Bonferroni.
data available:https://doi.org/10.25949/24295342

**Supplemental Table 5.** Mixed ANOVA *post hoc* comparisons for the interaction: ‘TMS condition’ x ‘carrier phrase length’.

| **Carrier Phrase** | **(I) TMS** | **(J) Sham** | **Mean Difference (I-J)** | **Std. Error** | **Sig.b** | **95% Confidence Interval for Difference b** |  |
|---|---|---|---|---|---|---|---|
|  |  |  |  |  |  | **Lower Bound** | **Upper Bound** |
| CP0 | 1.00 | 2.00 | -.139 | .197 | .488 | -.554 | .276 |
| CP1 | 1.00 | 2.00 | -.474<sup>*</sup> | .214 | .041 | -.926 | -.021 |
| CP2 | 1.00 | 2.00 | -.513<sup>*</sup> | .200 | .020 | -.935 | -.092 |
| CP3 | 1.00 | 2.00 | -.710<sup>*</sup> | .157 | <.001 | -1.042 | -.378 |

Based on estimated marginal means
\*. The mean difference is significant at the .05 level.
b. Adjustment for multiple comparisons: Bonferroni.
data available: https://doi.org/10.25949/24295342

**Supplemental Figure 1.**
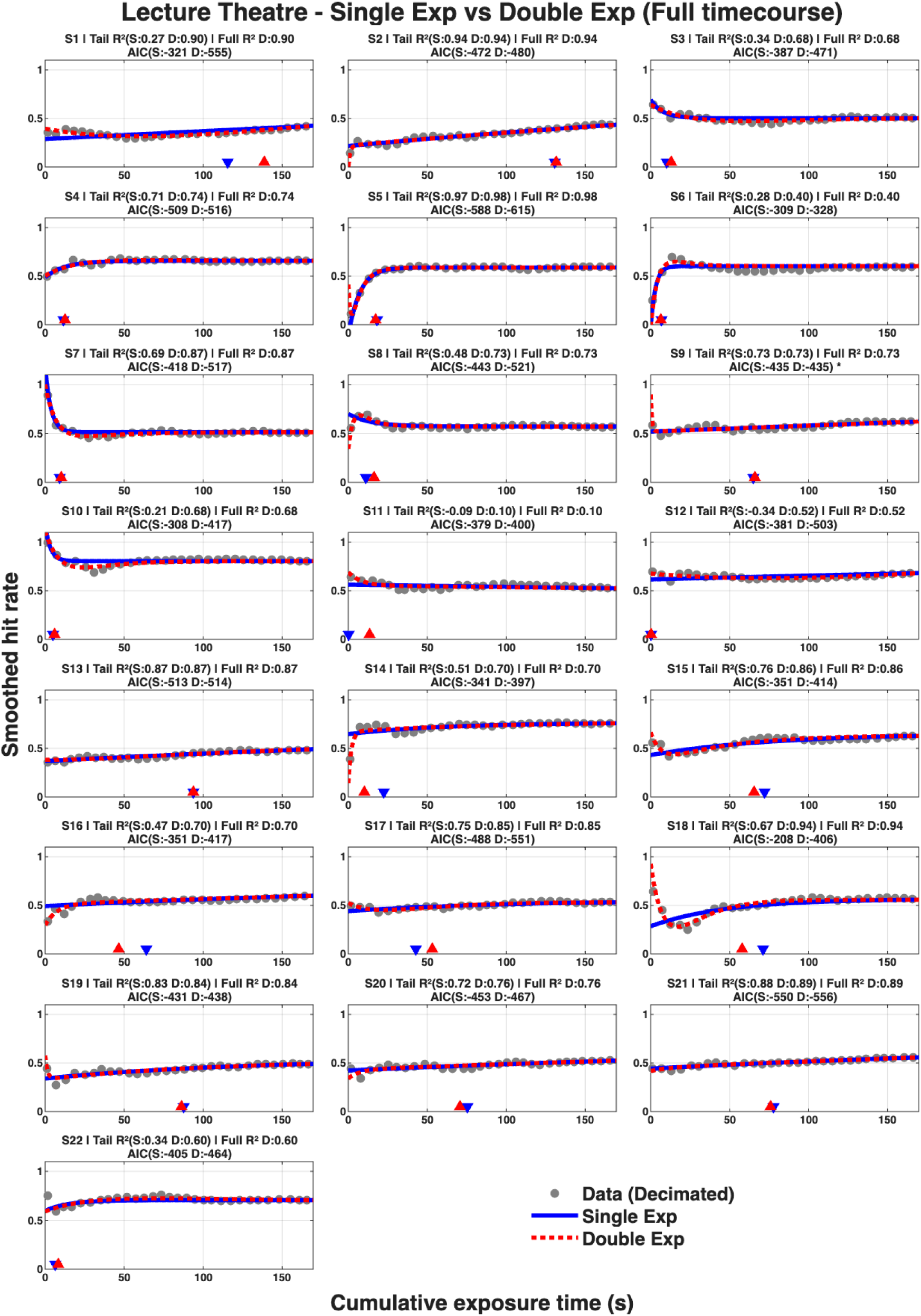
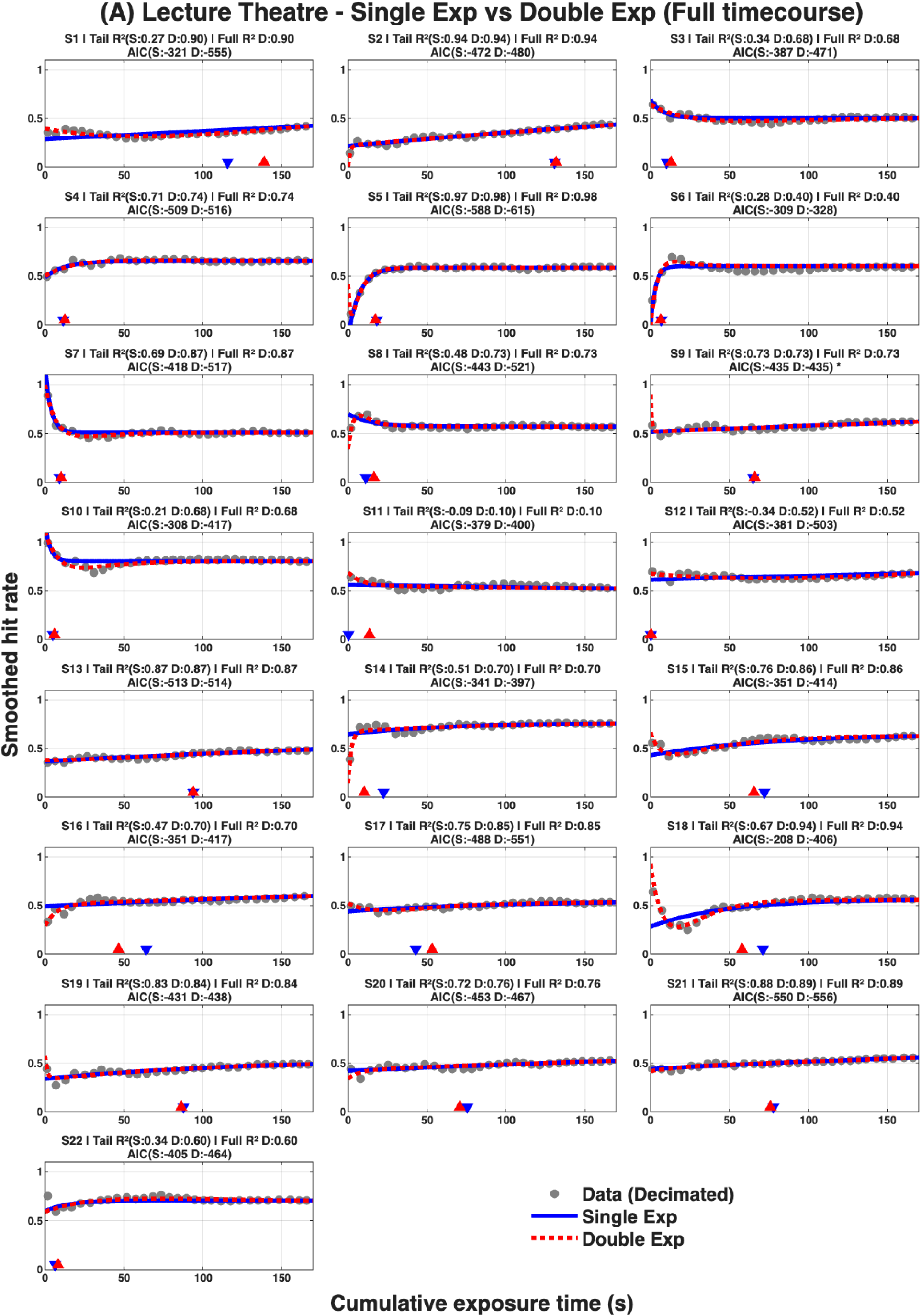
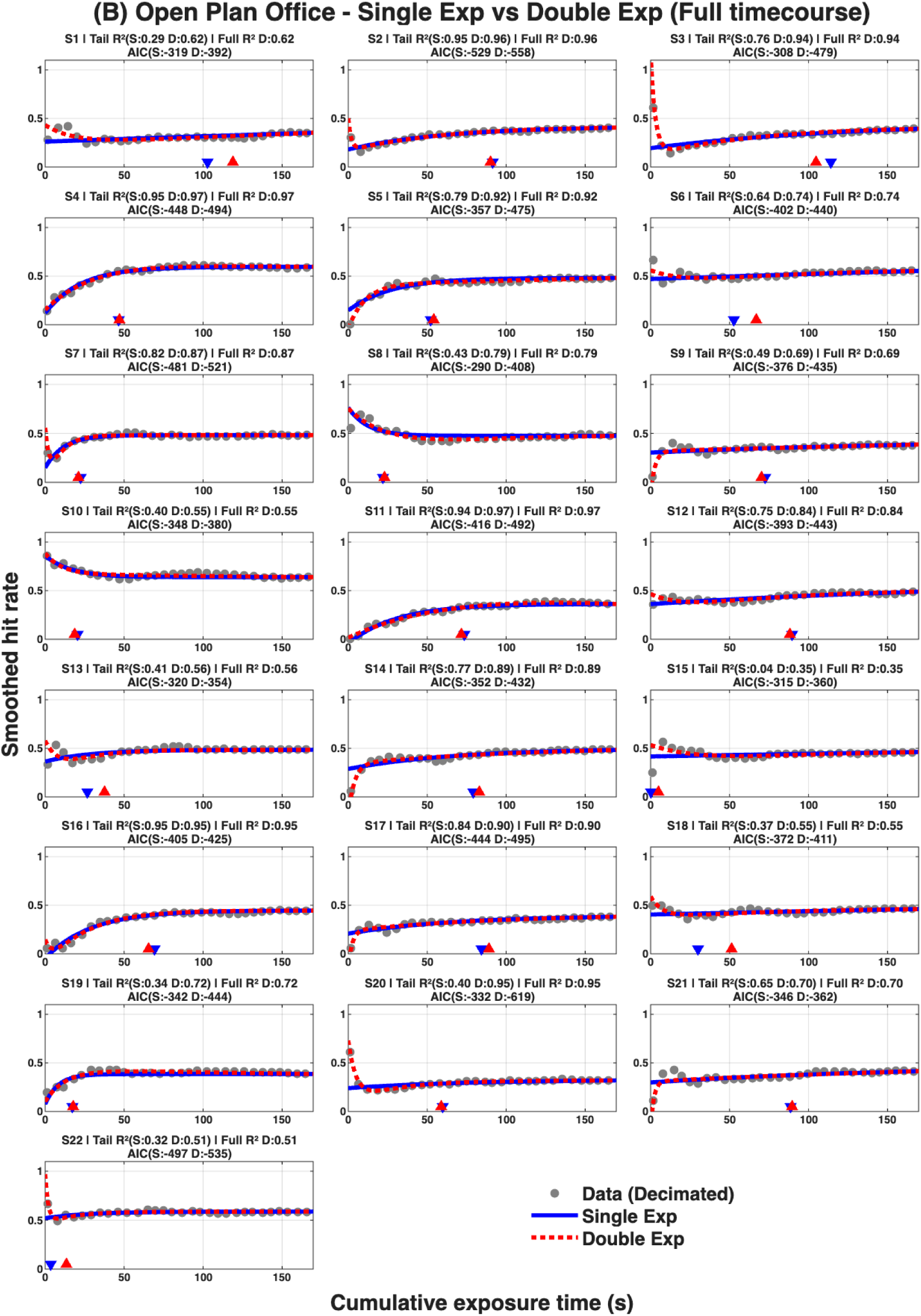
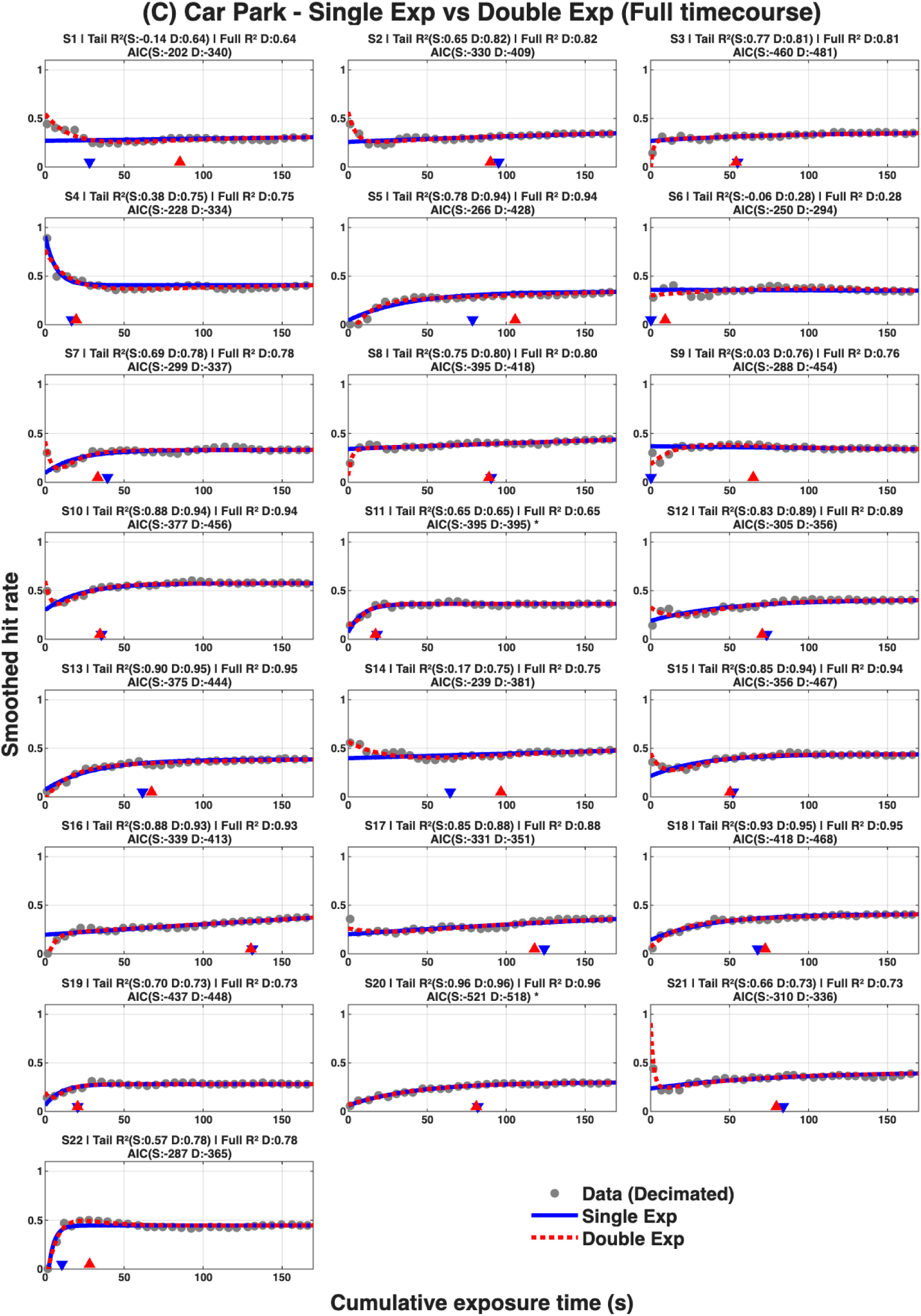
Time courses and single/double exponential fits of performance for 22 subjects in lecture theatre (A), open-plan office (B) and car park (C) where both exponential fits were performed using full data. Single (solid blue) and double (dotted red) exponential fits were fitted to the hit rate data (light grey marker, skipping every 4^th^ time point) that was smoothed using a 5-point moving average (equivalent to ∼7s). Akaike Information Criterion (AIC) and adjusted R^2^ (Tail R^2^) calculated on the truncated dataset are shown for both single (S) and double (D) exponential fits while full R^2^ is also quoted for the double exponential fit. All data available: https://doi.org/10.25949/24295342

**Supplemental Figure 2.**
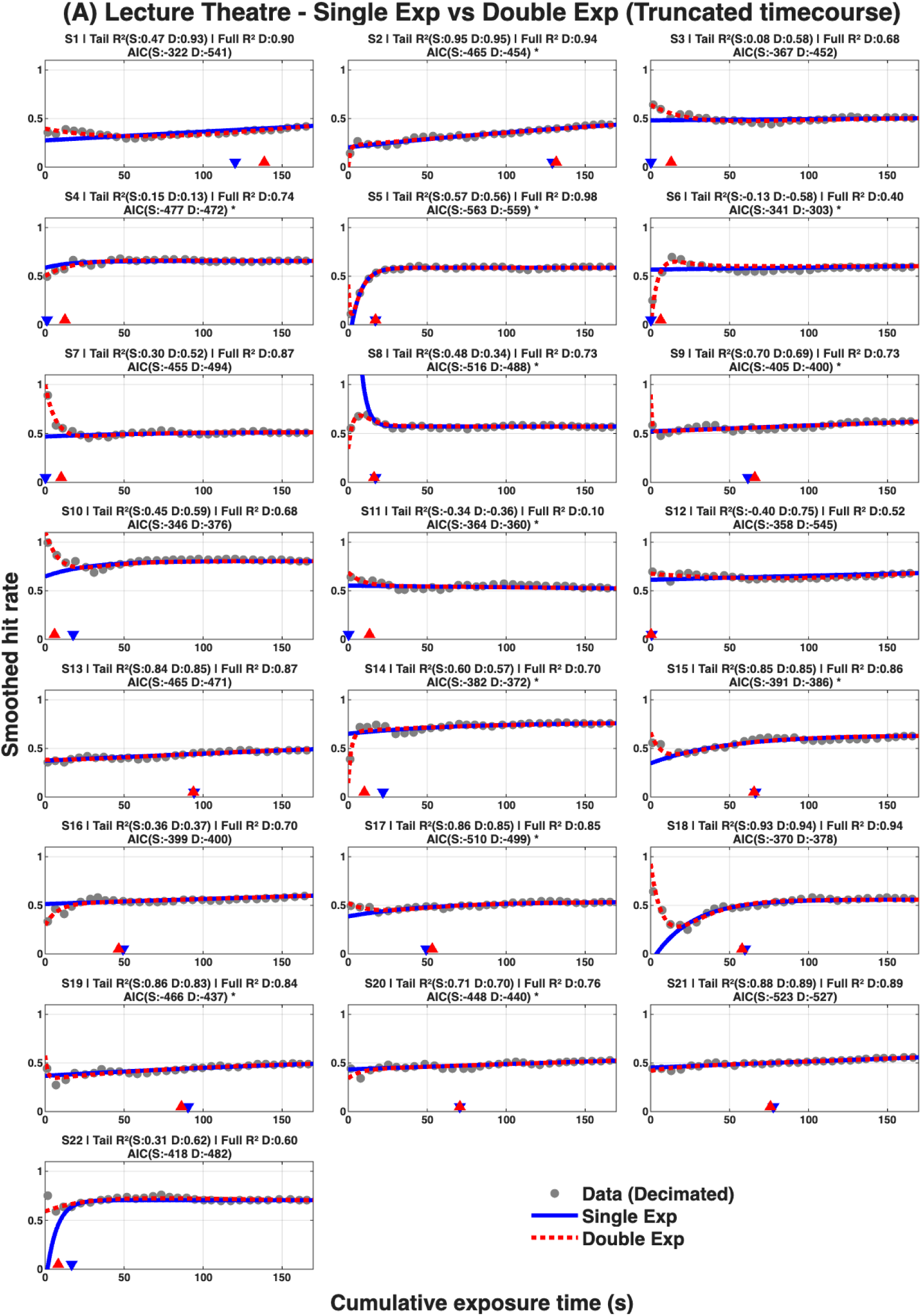
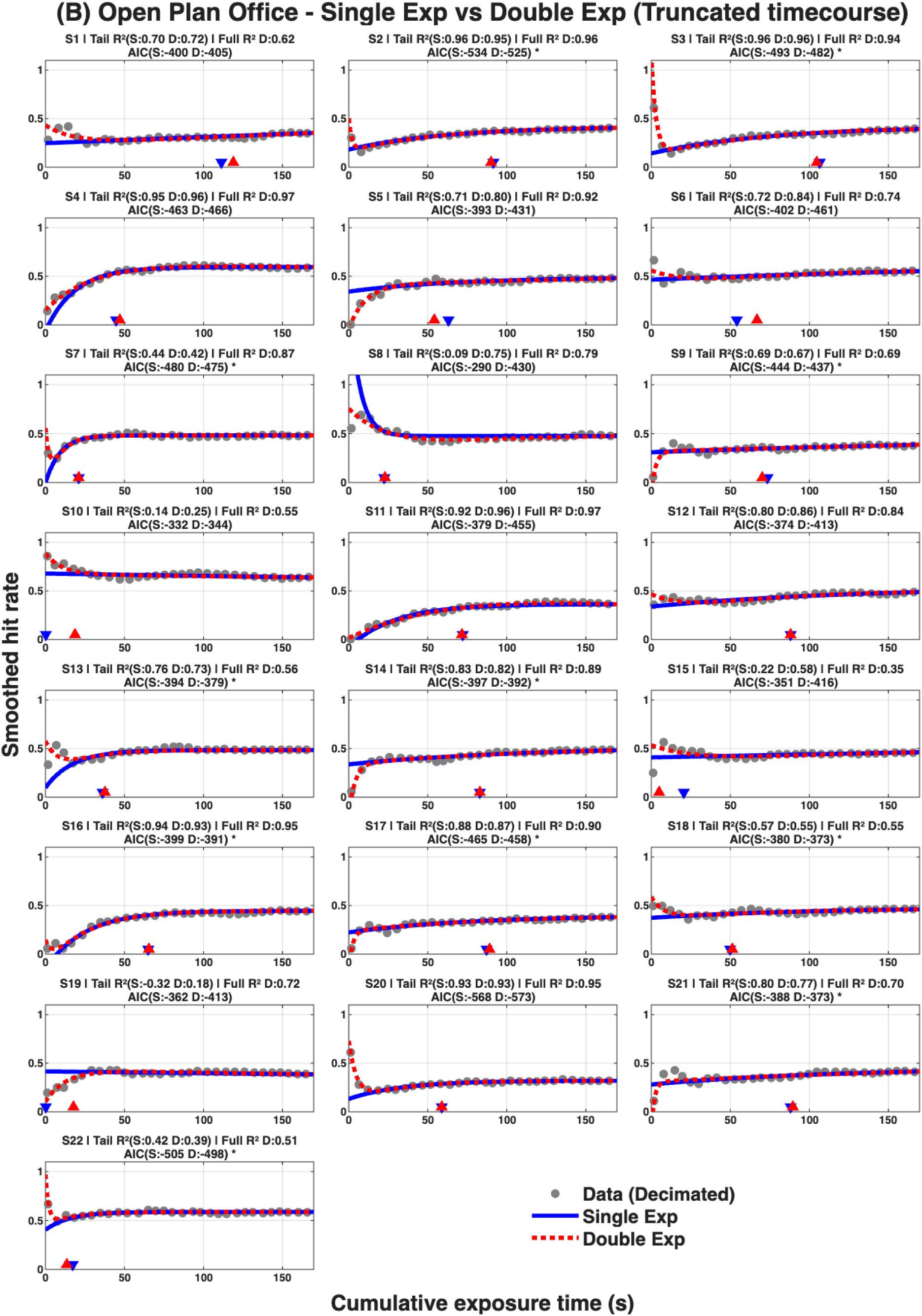
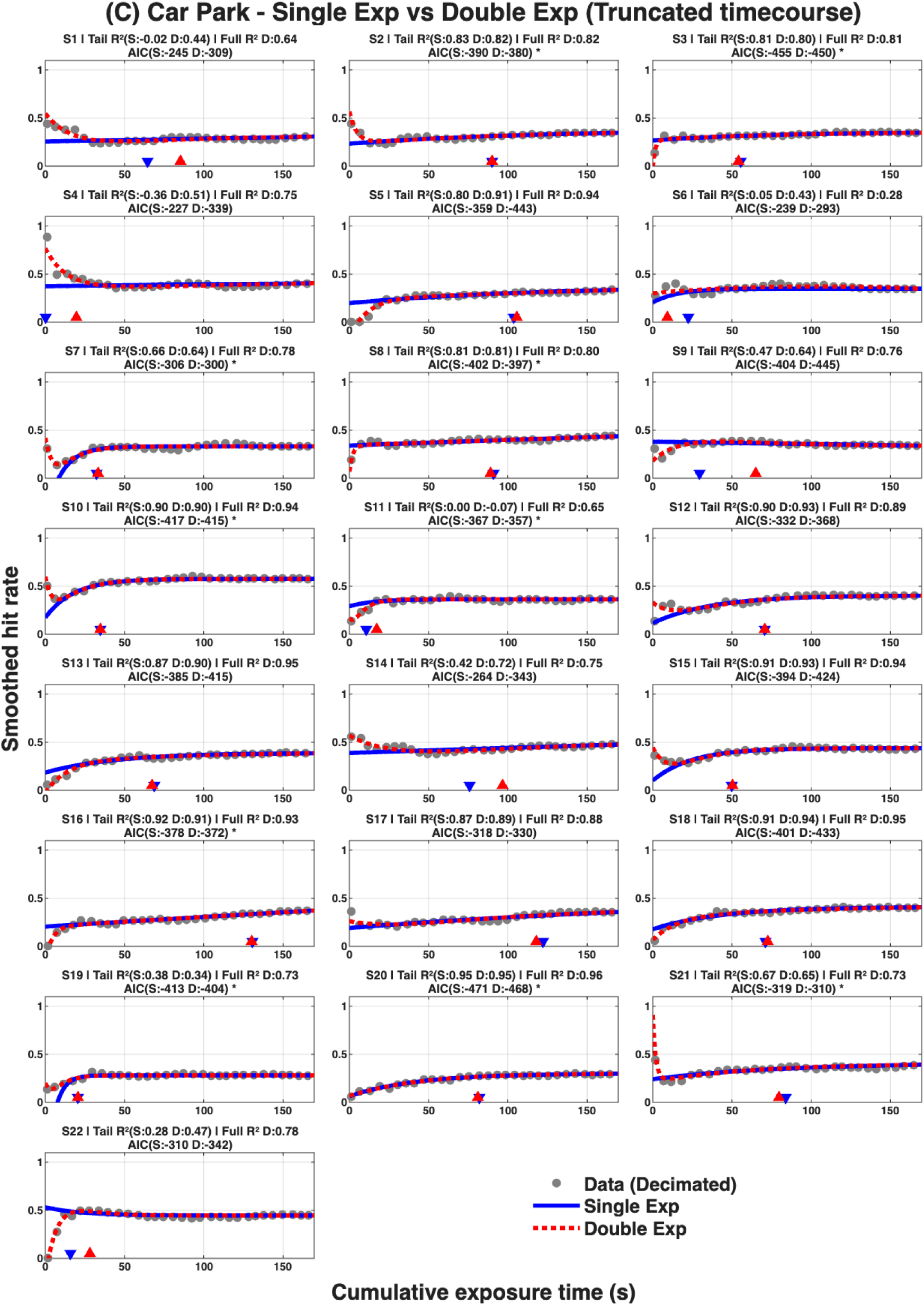
Time courses and single/double exponential fits of performance for 22 subjects in lecture theatre (A), open-plan office (B) and car park (C) where single exponential fits were calculated, ignoring the first 10 (∼14s) of data. Single (solid blue) and double (dotted red) exponential fits were fitted to the hit rate data (light grey marker, skipping every 4^th^ time point) that was smoothed using a 5-point moving average (equivalent to ∼7s). Akaike Information Criterion (AIC) and adjusted R^2^ (Tail R^2^) calculated on the truncated dataset are shown for both single (S) and double (D) exponential fits while full R^2^ is also quoted for the double exponential fit. All data available: https://doi.org/10.25949/24295342

**Supplemental Figure 3.**
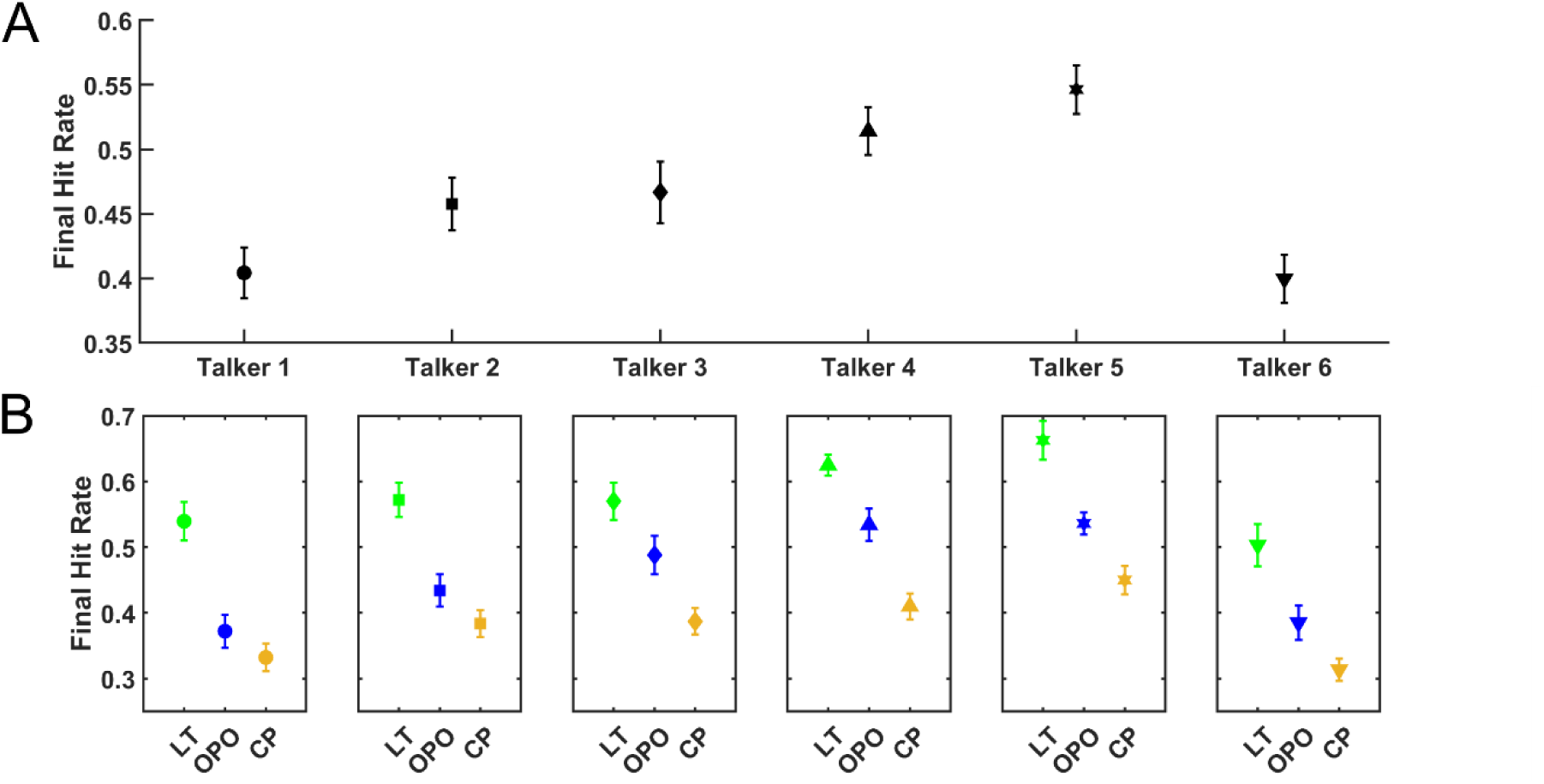
Comparison of Final Hit Rate (FHR) performances for different talkers. **A.** Performance, calculated as FHR in all rooms, is shown for the 6 different talkers (1-3 male and 4-6 female) presented to 22 individuals who did not undergo experimental TMS during the task involving the Lecture Room, Open-Plan Office and Car Park RIRs. **B.** Performance was also similarly calculated for the different talkers but is now separated by the room RIR associated with the specific talker. All data available: https://doi.org/10.25949/24295342

**Supplemental Figure 4.**
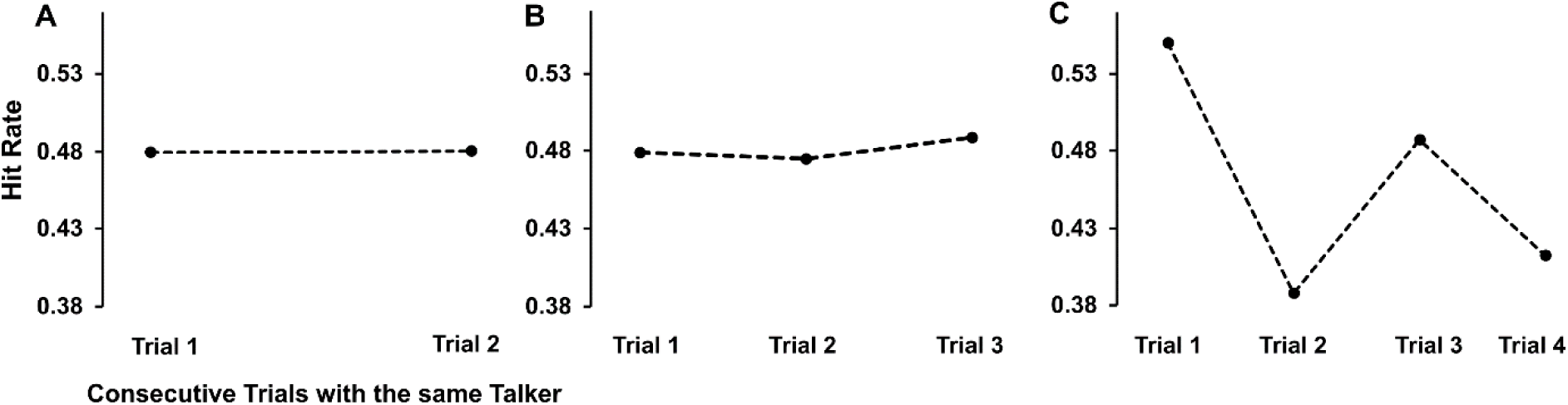
Hit Rate performances for |Color| and |Number| independent of environment presented or carrier phrase length. **A.** Performance, calculated for the 1262 trials where 22 participants heard in 2 consecutive trials the same talker, a Wilcoxon signed rank test revealed no significant differences in performance between trial 1 vs. trial 2 (n=1262, Z= −0.42, p=0.68, small effect size (r= 0.01)). **B.** Performance was also similarly calculated when listeners heard the same talker in three consecutive trials, this happened in a total of 217 trials and a Wilcoxon signed rank test revealed no significant differences in performance between trial 1 vs. trial 3 (n=217, Z= −0.17, p=0.87, small effect size (r= 0.01)). **C.** Performance calculated in 40 trials were listeners heard the same talker in four consecutive trials. No statistical differences were observed between Trial 1 and Trial 4 (n=40, Z= −0.159, p=0.11, small effect size (r= 0.02)). All data available: https://doi.org/10.25949/24295342

**Supplemental Figure 5.**
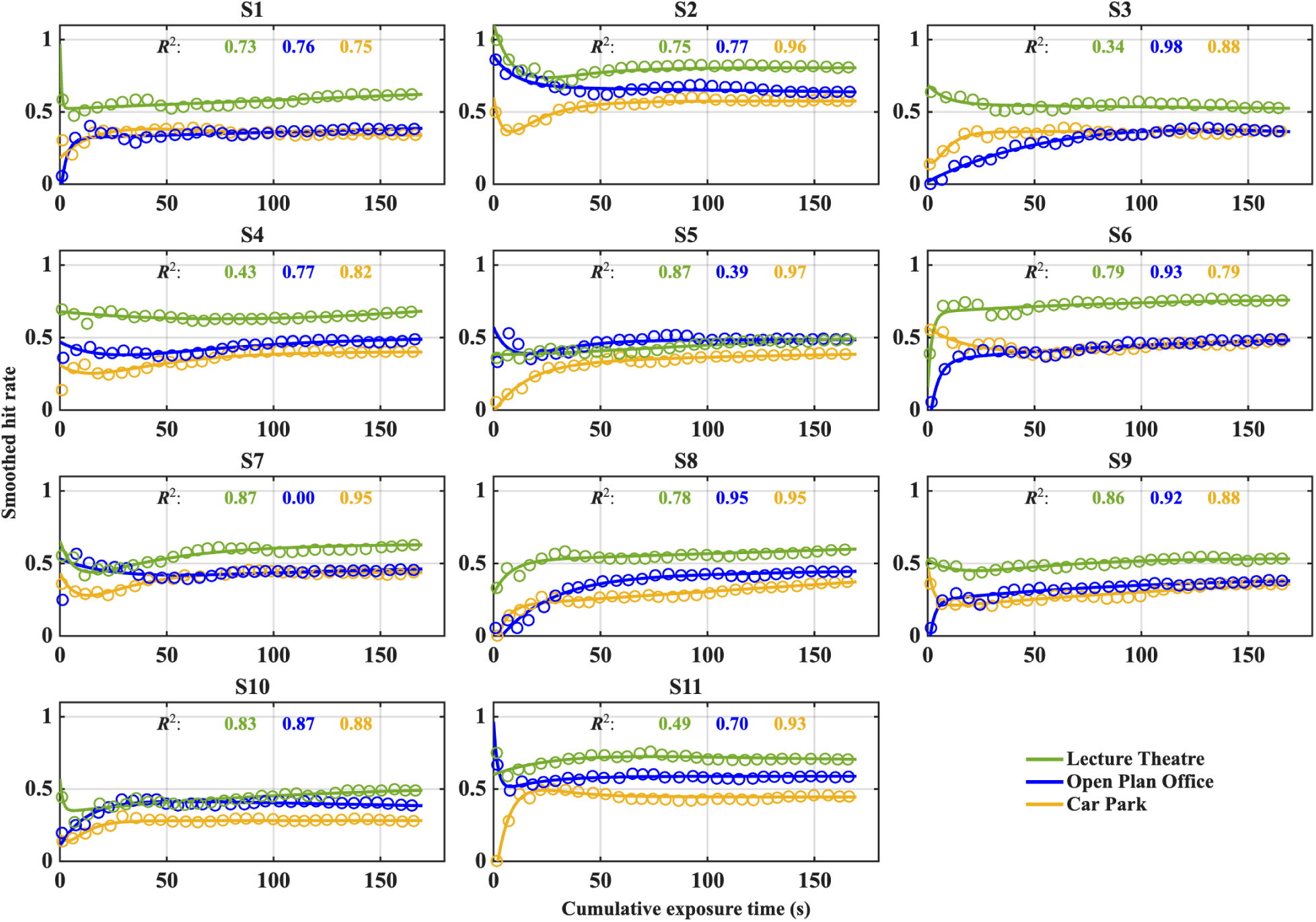
Time courses of performance for 11 sham-TMS subjects in lecture theatre (green), open plan office (blue) and car park (yellow). Solid lines represent actual cumulative hit rate data for each room that were calculated from raw data using a 5-point moving average (equivalent to ∼7s). Open circles represents best fit of double exponential with adjusted R2 values shown above for each individual. All data available: https://doi.org/10.25949/24295342

**Supplemental Figure 6.**
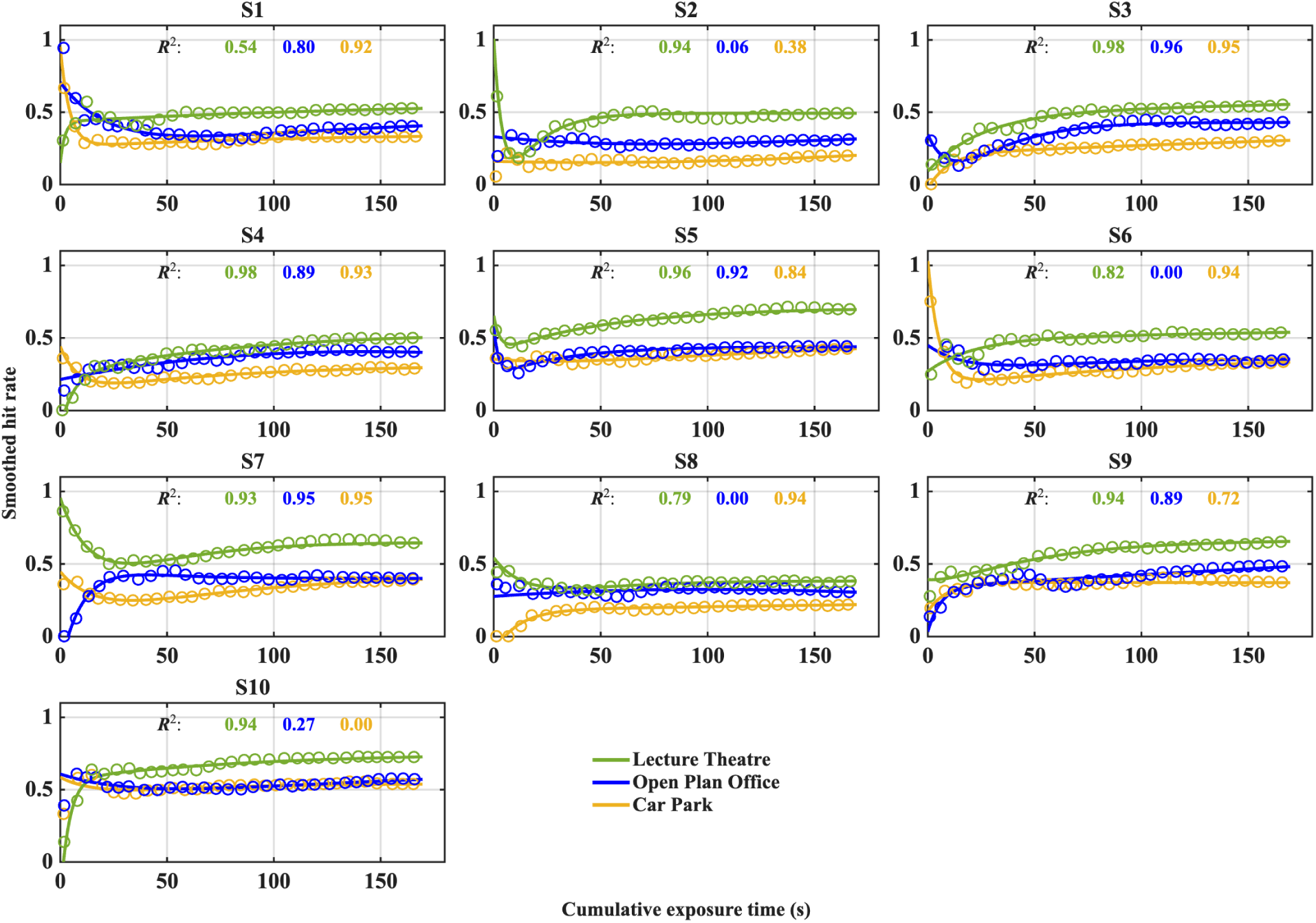
Time courses of performance for 10 TMS in lecture theatre (green), open plan office (blue) and car park (yellow). Solid lines represent actual cumulative hit rate data for each room that were calculated from raw data using a 5-point moving average (equivalent to ∼7s). Open circles represents best fit of double exponential with adjusted R2 values shown above for each individual. All data available: https://doi.org/10.25949/24295342

## Supplemental Information. Transcranial Magnetic Stimulation screening form

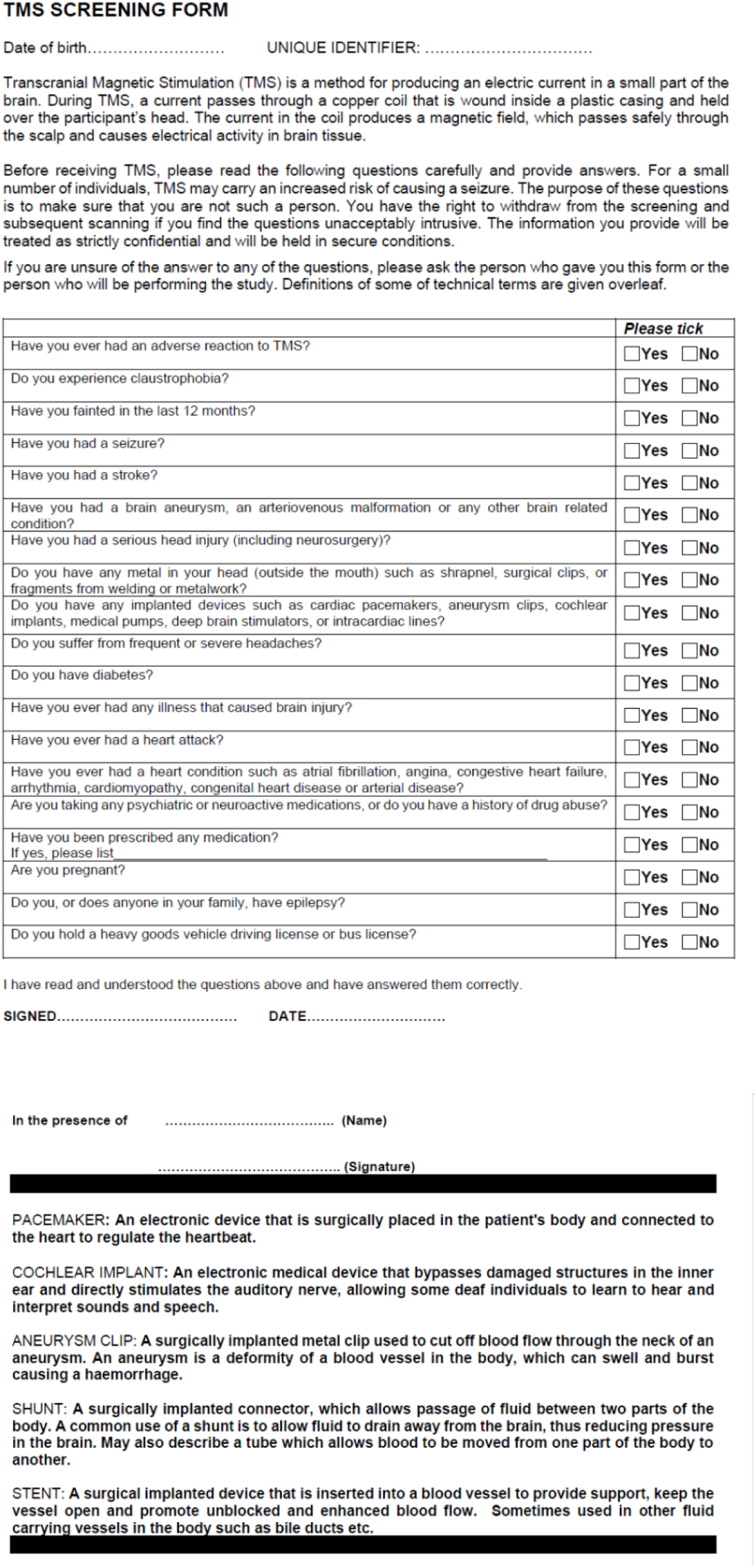

## Notes

https://doi.org/10.25949/24295342

### Competing Interest Statement

The authors have declared no competing interest.

### Summary of Updates

This manuscript was submitted to eLife and underwent peer review. Four reviewers assessed the original submission, and we revised the manuscript in response. This version (Revision 2) addresses the further comments received during the eLife review process. Reviewers 1 and 2 confirmed that all of their comments had been successfully addressed in Revision 1. Reviewers 3 and 4 provided further comments on that revision, which we address in this version. The eLife assessment and the reviewers public comments are publicly available. We have posted this preprint to share the work openly while peer review continues.

## References

Agus, T. R., Carrión-Castillo, A., Pressnitzer, D., & Ramus, F. (2014). Perceptual Learning of Acoustic Noise by Individuals With Dyslexia. Journal of Speech, Language, and Hearing Research: JSLHR, 57(3), 1069–1077. 10.1044/1092-4388(2013/13-0020)

Ainsworth, W. A., & Meyer, G. F. (1994). Recognition of plosive syllables in noise: comparison of an auditory model with human performance. The Journal of the Acoustical Society of America, 96(2 Pt 1), 687–694. 10.1121/1.410306

Albouy, G., Fogel, S., King, B. R., Laventure, S., Benali, H., Karni, A., Carrier, J., Robertson, E. M., & Doyon, J. (2015). Maintaining vs. enhancing motor sequence memories: respective roles of striatal and hippocampal systems. NeuroImage, 108, 423–434. 10.1016/j.neuroimage.2014.12.049

Alexander, W. H., & Brown, J. W. (2018). Frontal cortex function as derived from hierarchical predictive coding. Scientific Reports, 8(1), 3843. 10.1038/s41598-018-21407-9

Ambrus, G. G., Vékony, T., Janacsek, K., Trimborn, A. B. C., Kovács, G., & Nemeth, D. (2020). When less is more: Enhanced statistical learning of non-adjacent dependencies after disruption of bilateral DLPFC. Journal of Memory and Language, 114, 104144.

Anderson, L. A., & Malmierca, M. S. (2013). The effect of auditory cortex deactivation on stimulus-specific adaptation in the inferior colliculus of the rat. The European Journal of Neuroscience, 37(1), 52–62. 10.1111/ejn.12018

Antunes, F. M., & Malmierca, M. S. (2011). Effect of auditory cortex deactivation on stimulus-specific adaptation in the medial geniculate body. The Journal of Neuroscience: The Official Journal of the Society for Neuroscience, 31(47), 17306–17316. 10.1523/JNEUROSCI.1915-11.2011

Aslin, R. N., Saffran, J. R., & Newport, E. L. (1998). Computation of Conditional Probability Statistics by 8-Month-Old Infants. Psychological Science, 9(4), 321–324. 10.1111/1467-9280.00063

Badajoz-Davila, J., Buchholz, J. M., & Van-Hoesel, R. (2020). Effect of noise and reverberation on speech intelligibility for cochlear implant recipients in realistic sound environments. The Journal of the Acoustical Society of America, 147(5), 3538. 10.1121/10.0001259

Bajo, V. M., Nodal, F. R., Korn, C., Constantinescu, A. O., Mann, E. O., Boyden, E. S., 3rd, & King, A. J. (2019). Silencing cortical activity during sound-localization training impairs auditory perceptual learning. Nature Communications, 10(1), 3075. 10.1038/s41467-019-10770-4

Bakay, W. M. H., Anderson, L. A., Garcia-Lazaro, J. A., McAlpine, D., & Schaette, R. (2018). Hidden hearing loss selectively impairs neural adaptation to loud sound environments. Nature Communications, 9(1), 4298. 10.1038/s41467-018-06777-y

Barascud, N., Pearce, M. T., Griffiths, T. D., Friston, K. J., & Chait, M. (2016). Brain responses in humans reveal ideal observer-like sensitivity to complex acoustic patterns. Proceedings of the National Academy of Sciences, 113(5), E616–E625. 10.1073/pnas.1508523113

Barbas, H., & Pandya, D. N. (1987). Architecture and frontal cortical connections of the premotor cortex (area 6) in the rhesus monkey. The Journal of Comparative Neurology, 256(2), 211–228. 10.1002/cne.902560203

Barbas, H., & Pandya, D. N. (1991). Patterns of connections of the prefrontal cortex in the rhesus monkey associated with cortical architecture. In H. S. Levin (Ed.), Frontal lobe function and dysfunction, (pp (Vol. 427, pp. 35–58). Oxford University Press, xv. https://psycnet.apa.org/fulltext/1992-97203-002.pdf

Barberis, N. C. (2013). Thirty Years of Prospect Theory in Economics: A Review and Assessment. The Journal of Economic Perspectives: A Journal of the American Economic Association, 27(1), 173–196. 10.1257/jep.27.1.173

Bartolo, R., & Averbeck, B. B. (2021). Inference as a fundamental process in behavior. Current Opinion in Behavioral Sciences, 38, 8–13. 10.1016/j.cobeha.2020.06.005

Ben-David, B. M., Avivi-Reich, M., & Schneider, B. A. (2016). Does the degree of linguistic experience (native versus nonnative) modulate the degree to which listeners can benefit from a delay between the onset of the maskers and the onset of the target speech? Hearing Research, 341, 9–18. 10.1016/j.heares.2016.07.016

Bennett, I. J., Madden, D. J., Vaidya, C. J., Howard, J. H., Jr, & Howard, D. V. (2011). White matter integrity correlates of implicit sequence learning in healthy aging. Neurobiology of Aging, 32(12), 2317.e1-12. 10.1016/j.neurobiolaging.2010.03.017

Best, V., Ozmeral, E. J., Kopco, N., & Shinn-Cunningham, B. G. (2008). Object continuity enhances selective auditory attention. Proceedings of the National Academy of Sciences of the United States of America, 105(35), 13174–13178. 10.1073/pnas.0803718105

Bianco, R., Harrison, P. M. C., Hu, M., Bolger, C., Picken, S., Pearce, M. T., & Chait, M. (2020). Long-term implicit memory for sequential auditory patterns in humans. eLife, 9, e56073. 10.7554/eLife.56073

Binder, J. R., Liebenthal, E., Possing, E. T., Medler, D. A., & Ward, B. D. (2004). Neural correlates of sensory and decision processes in auditory object identification. Nature Neuroscience, 7(3), 295–301. 10.1038/nn1198

Blackwell, J. M., Lesicko, A. M., Rao, W., De Biasi, M., & Geffen, M. N. (2020). Auditory cortex shapes sound responses in the inferior colliculus. eLife, 9. 10.7554/eLife.51890

Blesser, B., & Salter, L.-R. (2009). Spaces Speak, Are You Listening?: Experiencing Aural Architecture. MIT Press. https://play.google.com/store/books/details?id=5aY1nrVTAZIC

Bolia, R. S., Nelson, W. T., Ericson, M. A., & Simpson, B. D. (2000). A speech corpus for multitalker communications research. The Journal of the Acoustical Society of America, 107(2), 1065–1066. 10.1121/1.428288

Bradley, J. S., Reich, R. D., & Norcross, S. G. (1999). On the combined effects of signal-to-noise ratio and room acoustics on speech intelligibility. The Journal of the Acoustical Society of America, 106(4 Pt 1), 1820–1828. 10.1121/1.427932

Brainard, M. S., & Doupe, A. J. (2002). What songbirds teach us about learning. Nature, 417(6886), 351–358. 10.1038/417351a

Brandewie, E. J., & Zahorik, P. (2018). Speech intelligibility in rooms: Disrupting the effect of prior listening exposure. The Journal of the Acoustical Society of America, 143(5), 3068. 10.1121/1.5038278

Brandewie, E., & Zahorik, P. (2010). Prior listening in rooms improves speech intelligibility. The Journal of the Acoustical Society of America, 128(1), 291–299. 10.1121/1.3436565

Brandewie, E., & Zahorik, P. (2011). Adaptation to Room Acoustics Using the Modified Rhyme Test. Proceedings of Meetings on Acoustics Acoustical Society of America, 129(4), 2487. 10.1121/1.3588198

Brandewie, E., & Zahorik, P. (2013). Time course of a perceptual enhancement effect for noise-masked speech in reverberant environments. The Journal of the Acoustical Society of America, 134(2), EL265-70. 10.1121/1.4816263

Bregman, A. S. (1994). Auditory Scene Analysis: The Perceptual Organization of Sound. MIT Press. https://play.google.com/store/books/details?id=jI8muSpAC5AC

Bronkhorst, A. W., & Houtgast, T. (1999). Auditory distance perception in rooms. Nature, 397(6719), 517–520. 10.1038/17374

Brumm, H., & Naguib, M. (2009). Chapter 1 Environmental Acoustics and the Evolution of Bird Song. In Advances in the Study of Behavior (Vol. 40, pp. 1–33). Academic Press. 10.1016/S0065-3454(09)40001-9

Brungart, D. S. (2001). Evaluation of speech intelligibility with the coordinate response measure. The Journal of the Acoustical Society of America, 109(5 Pt 1), 2276–2279. 10.1121/1.1357812

Cabrera, D., Jeong, C., Kwak, H. J., & Kim, J.-Y. (2005). Auditory room size perception for modeled and measured rooms. INTER-NOISE and NOISE-CON Congress and Conference Proceedings, 2005, 2995–3004. https://www.academia.edu/download/41945460/Auditory_room_size_perception_for_modele20160203-29142-1gyb08z.pdf

Castro-Meneses, L. J., Johnson, B. W., & Sowman, P. F. (2016). Vocal response inhibition is enhanced by anodal tDCS over the right prefrontal cortex. Experimental Brain Research. Experimentelle Hirnforschung. Experimentation Cerebrale, 234(1), 185–195. 10.1007/s00221-015-4452-0

Cherry, E. C. (1953). Some Experiments on the Recognition of Speech, with One and with Two Ears. The Journal of the Acoustical Society of America, 25(5), 975–979. 10.1121/1.1907229

Choi, J. Y., & Perrachione, T. K. (2019). Time and information in perceptual adaptation to speech. Cognition, 192, 103982. 10.1016/j.cognition.2019.05.019

Coltheart, M., Cox, R., Sowman, P., Morgan, H., Barnier, A., Langdon, R., Connaughton, E., Teichmann, L., Williams, N., & Polito, V. (2018). Belief, delusion, hypnosis, and the right dorsolateral prefrontal cortex: A transcranial magnetic stimulation study. Cortex; a Journal Devoted to the Study of the Nervous System and Behavior, 101, 234–248. 10.1016/j.cortex.2018.01.001

Conway, C. M. (2020). How does the brain learn environmental structure? Ten core principles for understanding the neurocognitive mechanisms of statistical learning. Neuroscience and Biobehavioral Reviews, 112, 279–299. 10.1016/j.neubiorev.2020.01.032

Conway, C. M., Bauernschmidt, A., Huang, S. S., & Pisoni, D. B. (2010). Implicit statistical learning in language processing: word predictability is the key. Cognition, 114(3), 356–371. 10.1016/j.cognition.2009.10.009

Conway, C. M., & Christiansen, M. H. (2001). Sequential learning in non-human primates. Trends in Cognitive Sciences, 5(12), 539–546. 10.1016/s1364-6613(00)01800-3

Conway, C. M., & Christiansen, M. H. (2005). Modality-constrained statistical learning of tactile, visual, and auditory sequences. Journal of Experimental Psychology. Learning, Memory, and Cognition, 31(1), 24–39. 10.1037/0278-7393.31.1.24

Cooke, M. (2006). A glimpsing model of speech perception in noise. The Journal of the Acoustical Society of America, 119(3), 1562–1573. 10.1121/1.2166600

Cooke, M., Scharenborg, O., & Meyer, B. T. (2022). The time course of adaptation to distorted speech. The Journal of the Acoustical Society of America, 151(4), 2636. 10.1121/10.0010235

Costalupes, J. A., Young, E. D., & Gibson, D. J. (1984). Effects of continuous noise backgrounds on rate response of auditory nerve fibers in cat. Journal of Neurophysiology, 51(6), 1326–1344. 10.1152/jn.1984.51.6.1326

Culling, J. F., Hodder, K. I., & Toh, C. Y. (2003). Effects of reverberation on perceptual segregation of competing voices. The Journal of the Acoustical Society of America, 114(5), 2871–2876. 10.1121/1.1616922

Daikhin, L., Raviv, O., & Ahissar, M. (2017). Auditory Stimulus Processing and Task Learning Are Adequate in Dyslexia, but Benefits From Regularities Are Reduced. Journal of Speech, Language, and Hearing Research: JSLHR, 60(2), 471–479. 10.1044/2016_JSLHR-H-16-0114

Daikoku, T., & Yumoto, M. (2023). Order of statistical learning depends on perceptive uncertainty. Current Research in Neurobiology, 4(100080), 100080. 10.1016/j.crneur.2023.100080

Davis, M. H., Ford, M. A., Kherif, F., & Johnsrude, I. S. (2011). Does semantic context benefit speech understanding through “top--down” processes? Evidence from time-resolved sparse fMRI. Journal of Cognitive Neuroscience, 23(12), 3914–3932. https://direct.mit.edu/jocn/article-abstract/23/12/3914/5277

Daw, N. D., Niv, Y., & Dayan, P. (2005). Uncertainty-based competition between prefrontal and dorsolateral striatal systems for behavioral control. Nature Neuroscience, 8(12), 1704–1711. 10.1038/nn1560

de Boer, J., & Thornton, A. R. D. (2008). Neural correlates of perceptual learning in the auditory brainstem: efferent activity predicts and reflects improvement at a speech-in-noise discrimination task. The Journal of Neuroscience: The Official Journal of the Society for Neuroscience, 28(19), 4929–4937.

Dean, I., Harper, N. S., & McAlpine, D. (2005). Neural population coding of sound level adapts to stimulus statistics. Nature Neuroscience, 8, 1684. 10.1038/nn1541

Dean, I., Robinson, B. L., Harper, N. S., & McAlpine, D. (2008). Rapid Neural Adaptation to Sound Level Statistics. The Journal of Neuroscience: The Official Journal of the Society for Neuroscience, 28(25), 6430–6438. 10.1523/jneurosci.0470-08.2008

Dienes, Z., & Hutton, S. (2013). Understanding hypnosis metacognitively: rTMS applied to left DLPFC increases hypnotic suggestibility. Cortex; a Journal Devoted to the Study of the Nervous System and Behavior, 49(2), 386–392. 10.1016/j.cortex.2012.07.009

Du, Y., Buchsbaum, B. R., Grady, C. L., & Alain, C. (2016). Increased activity in frontal motor cortex compensates impaired speech perception in older adults. Nature Communications, 7, 12241. 10.1038/ncomms12241

Faul, F., Erdfelder, E., Lang, A.-G., & Buchner, A. (2007). G*Power 3: a flexible statistical power analysis program for the social, behavioral, and biomedical sciences. Behavior Research Methods, 39(2), 175–191. 10.3758/bf03193146

Fiser, J., & Aslin, R. N. (2001). Unsupervised statistical learning of higher-order spatial structures from visual scenes. Psychological Science, 12(6), 499–504. 10.1111/1467-9280.00392

Fuglsang, S. A., Dau, T., & Hjortkjær, J. (2017). Noise-robust cortical tracking of attended speech in real-world acoustic scenes. NeuroImage, 156, 435–444. 10.1016/j.neuroimage.2017.04.026

Fuster, J. M., Bodner, M., & Kroger, J. K. (2000). Cross-modal and cross-temporal association in neurons of frontal cortex. Nature, 405(6784), 347–351. 10.1038/35012613

Gamboa, O. L., Antal, A., Moliadze, V., & Paulus, W. (2010). Simply longer is not better: reversal of theta burst after-effect with prolonged stimulation. Experimental Brain Research. Experimentelle Hirnforschung. Experimentation Cerebrale, 204(2), 181–187. 10.1007/s00221-010-2293-4

Gariépy, J.-F., Watson, K. K., Du, E., Xie, D. L., Erb, J., Amasino, D., & Platt, M. L. (2014). Social learning in humans and other animals. Frontiers in Neuroscience, 8, 58. 10.3389/fnins.2014.00058

Garinis, A. C., Glattke, T., & Cone, B. K. (2011). The MOC reflex during active listening to speech. Journal of Speech, Language, and Hearing Research: JSLHR, 54(5), 1464–1476.

Gibson, D. J., Young, E. D., & Costalupes, J. A. (1985). Similarity of dynamic range adjustment in auditory nerve and cochlear nuclei. Journal of Neurophysiology, 53(4), 940–958. 10.1152/jn.1985.53.4.940

Giraud, A. L., Garnier, S., Micheyl, C., Lina, G., Chays, A., & Chéry-Croze, S. (1997). Auditory efferents involved in speech-in-noise intelligibility. Neuroreport, 8(7), 1779–1783.

Glasberg, B. R., & Moore, B. C. (1990). Derivation of auditory filter shapes from notched-noise data. Hearing Research, 47(1–2), 103–138. 10.1016/0378-5955(90)90170-t

Goldman-Rakic, P. S., & Schwartz, M. L. (1982). Interdigitation of contralateral and ipsilateral columnar projections to frontal association cortex in primates. Science, 216(4547), 755–757. 10.1126/science.6177037

Gruters, K. G., Murphy, D. L. K., Jenson, C. D., Smith, D. W., Shera, C. A., & Groh, J. M. (2018). The eardrums move when the eyes move: A multisensory effect on the mechanics of hearing. Proceedings of the National Academy of Sciences of the United States of America, 115(6), E1309–E1318. 10.1073/pnas.1717948115

Hacker, M. J., & Ratcliff, R. (1979). A revised table of d’ for M-alternative forced choice. Perception & Psychophysics, 26(2), 168–170. 10.3758/bf03208311

Hackett, T. A., Stepniewska, I., & Kaas, J. H. (1999). Prefrontal connections of the parabelt auditory cortex in macaque monkeys. Brain Research, 817(1–2), 45–58. 10.1016/s0006-8993(98)01182-2

Hernández-Pérez, H., Mikiel-Hunter, J., McAlpine, D., Dhar, S., Boothalingam, S., Monaghan, J. J. M., & McMahon, C. M. (2021). Understanding degraded speech leads to perceptual gating of a brainstem reflex in human listeners. PLoS Biology, 19(10), e3001439. 10.1371/journal.pbio.3001439

Herwig, U., Satrapi, P., & Schönfeldt-Lecuona, C. (2003). Using the international 10-20 EEG system for positioning of transcranial magnetic stimulation. Brain Topography, 16(2), 95–99. https://link.springer.com/content/pdf/10.1023%2FB%3ABRAT.0000006333.93597.9d.pdf

Hicks, J. M., & McDermott, J. H. (2024). Noise schemas aid hearing in noise. Proceedings of the National Academy of Sciences of the United States of America, 121(47). 10.1073/pnas.2408995121

Hoogendam, J. M., Ramakers, G. M. J., & Di Lazzaro, V. (2010). Physiology of repetitive transcranial magnetic stimulation of the human brain. Brain Stimulation, 3(2), 95–118. 10.1016/j.brs.2009.10.005

Houtgast, T., & Steeneken, H. J. (1973). The modulation transfer function in room acoustics as a predictor of speech intelligibility. Acta Acustica United with Acustica, 28(1), 66–73. https://www.ingentaconnect.com/content/dav/aaua/1973/00000028/00000001/art00012

Houtgast, T., & Steeneken, H. J. M. (1985). A review of the MTF concept in room acoustics and its use for estimating speech intelligibility in auditoria. The Journal of the Acoustical Society of America, 77(3), 1069–1077. 10.1121/1.392224

Huang, Y.-Z., Edwards, M. J., Rounis, E., Bhatia, K. P., & Rothwell, J. C. (2005). Theta Burst Stimulation of the Human Motor Cortex. Neuron, 45(2), 201–206. 10.1016/j.neuron.2004.12.033

Hughson, W., & Westlake, H. (1944). Manual for program outline for rehabilitation of aural casualties both military and civilian. Transactions - American Academy of Ophthalmology and Otolaryngology. American Academy of Ophthalmology and Otolaryngology.

Huyck, J. J., & Wright, B. A. (2011). Late maturation of auditory perceptual learning. Developmental Science, 14(3), 614–621. 10.1111/j.1467-7687.2010.01009.x

Idemaru, K., & Holt, L. L. (2011). Word recognition reflects dimension-based statistical learning. Journal of Experimental Psychology. Human Perception and Performance, 37(6), 1939–1956. 10.1037/a0025641

Idemaru, K., & Holt, L. L. (2014). Specificity of dimension-based statistical learning in word recognition. Journal of Experimental Psychology. Human Perception and Performance, 40(3), 1009–1021. 10.1037/a0035269

Ivanov, A. Z., King, A. J., Willmore, B. D. B., Walker, K. M. M., & Harper, N. S. (2022). Cortical adaptation to sound reverberation. eLife, 11. 10.7554/eLife.75090

Jiang, Y., Chen, Y., Wei, L., Liu, F., Zheng, Z., Zhang, Z., Li, Z., Tang, Y., Wang, J., Xie, Q., Niu, C. M., & Zhu, C. (2025). Evaluation of scalp-based targeting methods of DLPFC for TMS therapy. Transcranial Magnetic Stimulation, 4(100095), 100095. 10.1016/j.transm.2025.100095

Jurcak, V., Okamoto, M., Singh, A., & Dan, I. (2005). Virtual 10–20 measurement on MR images for inter-modal linking of transcranial and tomographic neuroimaging methods. NeuroImage, 26(4), 1184–1192. https://ac.els-cdn.com/S1053811905001862/1-s2.0-S1053811905001862-main.pdf?_tid=87d3462c-b7fa4780-b7b0-289d930895bd&acdnat=1544065940_494439b3ca37d9077258347769daf7e0

Kakehi, K. (1992). Adaptability to differences between talkers in Japanese monosyllabic perception. Speech Perception, Speech Production, and Linguistic Structure, 135–142. https://books.google.com.au/books?hl=en&lr=&id=eYrOR7VHnnQC&oi=fnd&pg=PA135&dq=Kakehi,+K.+(1992).+%E2%80%9CAdaptability+to+differences+between+talkers+in+Japanese+monosyllabic+perception,%E2%80%9D+in+Speech+Perception,+Speech+Production+and+Linguistic+Structure,+edited+by+Y.+Tohkura,+E.+Vatikiotis-Bateson,+and+Y.+Sagisaka+(OHM,+Tokyo),+pp.+135&ots=p9p7gXYBWu&sig=GPeZemflBK64RT6sNEwoGteaYXw

Kato, K., & Kakehi, K. (1988). Listener adaptability to individual speaker differences in monosyllabic speech perception. J. Acoust. Soc. Jpn, 44(3), 180–186.

Kell, A. J. E., & McDermott, J. H. (2019). Invariance to background noise as a signature of non-primary auditory cortex. Nature Communications, 10(1), 3958. 10.1038/s41467-019-11710-y

Khalighinejad, B., Herrero, J. L., Mehta, A. D., & Mesgarani, N. (2019). Adaptation of the human auditory cortex to changing background noise. Nature Communications, 10(1), 2509. 10.1038/s41467-019-10611-4

Kirkham, N. Z., Slemmer, J. A., & Johnson, S. P. (2002). Visual statistical learning in infancy: evidence for a domain general learning mechanism. Cognition, 83(2), B35–42. 10.1016/s0010-0277(02)00004-5

Knudsen, V. O. (1929). THE HEARING OF SPEECH IN AUDITORIUMS. The Journal of the Acoustical Society of America, 1(1), 56–82. 10.1121/1.1901470

Kolarik, A. J., Moore, B. C. J., Cirstea, S., Aggius-Vella, E., Gori, M., Campus, C., & Pardhan, S. (2021). Factors Affecting Auditory Estimates of Virtual Room Size: Effects of Stimulus, Level, and Reverberation. Perception, 50(7), 646–663. 10.1177/03010066211020598

Kressner, A. A., Westermann, A., & Buchholz, J. M. (2018). The impact of reverberation on speech intelligibility in cochlear implant recipients. The Journal of the Acoustical Society of America, 144(2), 1113. 10.1121/1.5051640

Larsby, B., Hällgren, M., Nilsson, L., & McAllister, A. (2015). The influence of female versus male speakers’ voice on speech recognition thresholds in noise: Effects of low- and high-frequency hearing impairment. Speech, Language and Hearing, 18, 83–90. 10.1179/2050572814Y.0000000053

Lauay, C., Gerlach, N. M., Adkins-Regan, E., & DeVoogd, T. J. (2004). Female zebra finches require early song exposure to prefer high-quality song as adults. Animal Behaviour, 68(6), 1249–1255. 10.1016/j.anbehav.2003.12.025

Lewicki, M. S., Olshausen, B. A., Surlykke, A., & Moss, C. F. (2014). Scene analysis in the natural environment. Frontiers in Psychology, 5, 199. 10.3389/fpsyg.2014.00199

Liao, L.-D., Tsytsarev, V., Delgado-Martínez, I., Li, M.-L., Erzurumlu, R., Vipin, A., Orellana, J., Lin, Y.-R., Lai, H.-Y., Chen, Y.-Y., & Thakor, N. V. (2013). Neurovascular coupling: in vivo optical techniques for functional brain imaging. Biomedical Engineering Online, 12(1), 38. 10.1186/1475-925X-12-38

Liu, R., & Holt, L. L. (2015). Dimension-based statistical learning of vowels. Journal of Experimental Psychology. Human Perception and Performance, 41(6), 1783–1798. 10.1037/xhp0000092

Lochner, J. P. A., & Burger, J. F. (1961). The intelligibility of speech under reverberant conditions. Acta Acustica United with Acustica, 11(4), 195–200. https://www.ingentaconnect.com/content/dav/aaua/1961/00000011/00000004/art00004

Marler, P. (1970). A comparative approach to vocal learning: Song development in white-crowned sparrows. Journal of Comparative and Physiological Psychology, 71(2p2), 1–25. 10.1037/h0029144

Mathews, R. C., Buss, R. R., Stanley, W. B., Blanchard-Fields, F., Cho, J. R., & Druhan, B. (1989). Role of implicit and explicit processes in learning from examples: A synergistic effect. Journal of Experimental Psychology. Learning, Memory, and Cognition, 15(6), 1083–1100. 10.1037/0278-7393.15.6.1083

McAlpine, D., & de Hoz, L. (2023). Listening loops and the adapting auditory brain. Frontiers in Neuroscience, 17, 1081295. 10.3389/fnins.2023.1081295

McDermott, J. H., Schemitsch, M., & Simoncelli, E. P. (2013). Summary statistics in auditory perception. Nature Neuroscience, 16(4), 493–498.

McDermott, J. H., & Simoncelli, E. P. (2011). Sound texture perception via statistics of the auditory periphery: evidence from sound synthesis. Neuron, 71(5), 926–940.

McNutt, M. K., Bradford, M., Drazen, J. M., Hanson, B., Howard, B., Jamieson, K. H., Kiermer, V., Marcus, E., Pope, B. K., Schekman, R., Swaminathan, S., Stang, P. J., & Verma, I. M. (2018). Transparency in authors’ contributions and responsibilities to promote integrity in scientific publication. Proceedings of the National Academy of Sciences of the United States of America, 115(11), 2557–2560. 10.1073/pnas.1715374115

McWalter, R., & McDermott, J. H. (2018). Adaptive and Selective Time Averaging of Auditory Scenes. Current Biology: CB, 28(9), 1405–1418.e10. 10.1016/j.cub.2018.03.049

Mesgarani, N., David, S. V., Fritz, J. B., & Shamma, S. A. (2014). Mechanisms of noise robust representation of speech in primary auditory cortex. Proceedings of the National Academy of Sciences of the United States of America, 111(18), 6792–6797. 10.1073/pnas.1318017111

Mishra, S. K., & Lutman, M. E. (2014). Top-down influences of the medial olivocochlear efferent system in speech perception in noise. PloS One, 9(1), e85756.

Morrone, M. C. (2010). Brain development: critical periods for cross-sensory plasticity. Current Biology: CB, 20(21), R934–6. 10.1016/j.cub.2010.09.052

Nielsen, J. B., & Dau, T. (2010). Revisiting perceptual compensation for effects of reverberation in speech identification. The Journal of the Acoustical Society of America, 128(5), 3088–3094. 10.1121/1.3494508

Nikolin, S., D’Souza, O., Vulovic, V., Alonzo, A., Chand, N., Dong, V., Martin, D., & Loo, C. (2019). Comparison of Site Localization Techniques for Brain Stimulation. The Journal of ECT, 35(2). https://journals.lww.com/ectjournal/fulltext/2019/06000/comparison_of_site_localization_techniques_for.14.aspx

Nissen, M. J., & Bullemer, P. (1987). Attentional requirements of learning: Evidence from performance measures. Cognitive Psychology, 19(1), 1–32. 10.1016/0010-0285(87)90002-8

Nydam, A. S., Sewell, D. K., & Dux, P. E. (2018). Cathodal electrical stimulation of frontoparietal cortex disrupts statistical learning of visual configural information. Cortex; a Journal Devoted to the Study of the Nervous System and Behavior, 99, 187–199. 10.1016/j.cortex.2017.11.008

Packard, M. G., & Knowlton, B. J. (2002). Learning and memory functions of the Basal Ganglia. Annual Review of Neuroscience, 25(1), 563–593. 10.1146/annurev.neuro.25.112701.142937

Pallant, J. (2011). A Step by Step Guide to Data Analysis Using SPSS. https://www.academia.edu/download/38306978/_Julie_Pallant_SPSS_Survival_Manual_A_Step_by_St1.pdf

Pandya, D. Ν., & Barnes, C. L. (2019). Architecture and connections of the frontal lobe. The Frontal Lobes Revisited. 10.4324/9781315788975-3/architecture-connections-frontal-lobe-deepak-pandya-clifford-barnes

Pascual-Leone, A., Wassermann, E. M., Grafman, J., & Hallett, M. (1996). The role of the dorsolateral prefrontal cortex in implicit procedural learning. Experimental Brain Research. Experimentelle Hirnforschung. Experimentation Cerebrale, 107(3), 479–485. 10.1007/BF00230427

Perrot, X., Ryvlin, P., Isnard, J., Guénot, M., Catenoix, H., Fischer, C., Mauguière, F., & Collet, L. (2006). Evidence for corticofugal modulation of peripheral auditory activity in humans. Cerebral Cortex, 16(7), 941–948.

Petrides, M., & Pandya, D. N. (2002). Comparative cytoarchitectonic analysis of the human and the macaque ventrolateral prefrontal cortex and corticocortical connection patterns in the monkey. The European Journal of Neuroscience, 16(2), 291–310. 10.1046/j.1460-9568.2001.02090.x

Phillips, D. P. (1985). Temporal response features of cat auditory cortex neurons contributing to sensitivity to tones delivered in the presence of continuous noise. Hearing Research, 19(3), 253–268. 10.1016/0378-5955(85)90145-5

Plakke, B., & Romanski, L. M. (2014). Auditory connections and functions of prefrontal cortex. Frontiers in Neuroscience, 8, 199. 10.3389/fnins.2014.00199

Reber, A. S. (1967). Implicit learning of artificial grammars. Journal of Verbal Learning and Verbal Behavior, 6(6), 855–863. 10.1016/S0022-5371(67)80149-X

Rebuschat, P. (2015). Implicit and Explicit Learning of Languages. John Benjamins Publishing Company. https://play.google.com/store/books/details?id=jI-ACgAAQBAJ

Rees, A., & Palmer, A. R. (1988). Rate-intensity functions and their modification by broadband noise for neurons in the guinea pig inferior colliculus. The Journal of the Acoustical Society of America. https://pubs.aip.org/asa/jasa/article-abstract/83/4/1488/826235

Robinson, B. L., Harper, N. S., & McAlpine, D. (2016). Meta-adaptation in the auditory midbrain under cortical influence. Nature Communications, 7, 13442. 10.1038/ncomms13442

Romero, M. C., Merken, L., Janssen, P., & Davare, M. (2022). Neural effects of continuous theta-burst stimulation in macaque parietal neurons. eLife, 11, e65536. 10.7554/eLife.65536

Sabine, H. (1953). Room acoustics. Trans IRE, 1, 4–12. https://www.theatrecrafts.com/archive/cue/cue_18_5.pdf

Saffran, J. R. (2003). Statistical Language Learning: Mechanisms and Constraints. Current Directions in Psychological Science, 12(4), 110–114. 10.1111/1467-8721.01243

Saffran, J. R., Aslin, R. N., & Newport, E. L. (1996). Statistical learning by 8-month-old infants. Science, 274(5294), 1926–1928. 10.1126/science.274.5294.1926

Saffran, J. R., Johnson, E. K., Aslin, R. N., & Newport, E. L. (1999). Statistical learning of tone sequences by human infants and adults. Cognition, 70(1), 27–52. 10.1016/s0010-0277(98)00075-4

Salvi, R. J., Lockwood, A. H., Frisina, R. D., Coad, M. L., Wack, D. S., & Frisina, D. R. (2002). PET imaging of the normal human auditory system: responses to speech in quiet and in background noise. Hearing Research, 170(1–2), 96–106. 10.1016/s0378-5955(02)00386-6

Santon, F. (1976). Numerical prediction of echograms and of the intelligibility of speech in rooms. The Journal of the Acoustical Society of America, 59(6), 1399–1405. 10.1121/1.381027

Schroeder, M. R. (1962). Frequency-Correlation Functions of Frequency Responses in Rooms. The Journal of the Acoustical Society of America, 34(12), 1819–1823. 10.1121/1.1909136

Schultz, W. (2002). Getting formal with dopamine and reward. Neuron, 36(2), 241–263. 10.1016/s0896-6273(02)00967-4

Shapiro, S., & Wilk, M. (1965). An analysis of variance test for normality (complete samples). Biometrika, 52(3–4), 591–611. 10.1093/BIOMET/52.3-4.591

Shinn-Cunningham, B. (2000, July). Learning Reverberation: Considerations for Spatial Auditory Displays. https://www.researchgate.net/profile/Barbara-Shinn-Cunningham/publication/2414695_Learning_Reverberation_Considerations_for_Spatial_Auditory_Displays/links/0f31752f91fb83f924000000/Learning-Reverberation-Considerations-for-Spatial-Auditory-Displays.pdf

Shinn-Cunningham, B., & Kawakyu, K. (2003). Neural representation of source direction in reverberant space. 2003 IEEE Workshop on Applications of Signal Processing to Audio and Acoustics (IEEE Cat. No.03TH8684), 79–82. 10.1109/ASPAA.2003.1285824

Simpson, A. J. R., Harper, N. S., Reiss, J. D., & McAlpine, D. (2014). Selective adaptation to “oddball” sounds by the human auditory system. The Journal of Neuroscience: The Official Journal of the Society for Neuroscience, 34(5), 1963–1969. 10.1523/JNEUROSCI.4274-13.2013

Smith, E. C., & Lewicki, M. S. (2006). Efficient auditory coding. Nature, 439(7079), 978–982. 10.1038/nature04485

Srinivasan, N. K., & Zahorik, P. (2012). Prior listening exposure to a reverberant room improves open-set intelligibility of high-variability sentences. The Journal of the Acoustical Society of America, 133(1), EL33–EL39. 10.1121/1.4771978

Stillman, C. M., Gordon, E. M., Simon, J. R., Vaidya, C. J., Howard, D. V., & Howard, J. H., Jr. (2013). Caudate resting connectivity predicts implicit probabilistic sequence learning. Brain Connectivity, 3(6), 601–610. 10.1089/brain.2013.0169

Stilp, C. (2020). Acoustic context effects in speech perception. Wiley Interdisciplinary Reviews. Cognitive Science, 11(1), e1517. 10.1002/wcs.1517

Takács, Á., Kóbor, A., Kardos, Z., Janacsek, K., Horváth, K., Beste, C., & Nemeth, D. (2021). Neurophysiological and functional neuroanatomical coding of statistical and deterministic rule information during sequence learning. Human Brain Mapping, 42(10), 3182–3201. 10.1002/hbm.25427

Taylor, S. F., Gu, P., Simmonite, M., Lasagna, C., Tso, I. F., Lee, T. G., Vesia, M., & Hernandez-Garcia, L. (2024). Lateral prefrontal stimulation of active cortex with theta burst transcranial magnetic stimulation affects subsequent engagement of the frontoparietal network. Biological Psychiatry: Cognitive Neuroscience and Neuroimaging, 9(2), 235–244. 10.1016/j.bpsc.2023.10.005

Terreros, G., & Delano, P. H. (2015). Corticofugal modulation of peripheral auditory responses. Frontiers in Systems Neuroscience, 9. 10.3389/fnsys.2015.00134

Traer, J., & McDermott, J. H. (2016). Statistics of natural reverberation enable perceptual separation of sound and space. Proceedings of the National Academy of Sciences, 113(48), E7856–E7865. 10.1073/pnas.1612524113

Trapp, N. T., Bruss, J., King Johnson, M., Uitermarkt, B. D., Garrett, L., Heinzerling, A., Wu, C., Koscik, T. R., Ten Eyck, P., & Boes, A. D. (2020). Reliability of targeting methods in TMS for depression: Beam F3 vs. 5.5 cm. Brain Stimulation, 13(3), 578–581. 10.1016/j.brs.2020.01.010

Tupak, S. V., Dresler, T., Badewien, M., Hahn, T., Ernst, L. H., Herrmann, M. J., Deckert, J., Ehlis, A.-C., & Fallgatter, A. J. (2013). Inhibitory transcranial magnetic theta burst stimulation attenuates prefrontal cortex oxygenation. Human Brain Mapping, 34(1), 150–157. 10.1002/hbm.21421

van den Bos, R., Jolles, J. W., & Homberg, J. R. (2013). Social modulation of decision-making: a cross-species review. Frontiers in Human Neuroscience, 7, 301. 10.3389/fnhum.2013.00301

Varnava, A., Stokes, M. G., & Chambers, C. D. (2011). Reliability of the “observation of movement” method for determining motor threshold using transcranial magnetic stimulation. Journal of Neuroscience Methods, 201(2), 327–332. 10.1016/j.jneumeth.2011.08.016

Vékony, T., Ambrus, G. G., Janacsek, K., & Nemeth, D. (2022). Cautious or causal? Key implicit sequence learning paradigms should not be overlooked when assessing the role of DLPFC (Commentary on Prutean et al.). Cortex; a Journal Devoted to the Study of the Nervous System and Behavior, 148, 222–226. 10.1016/j.cortex.2021.10.001

Vlahou, E., Seitz, A. R., & Kopčo, N. (2019). Nonnative implicit phonetic training in multiple reverberant environments. Attention, Perception & Psychophysics, 81(4), 935–947. 10.3758/s13414-019-01680-0

Wächter, T., Lungu, O. V., Liu, T., Willingham, D. T., & Ashe, J. (2009). Differential effect of reward and punishment on procedural learning. The Journal of Neuroscience: The Official Journal of the Society for Neuroscience, 29(2), 436–443. 10.1523/JNEUROSCI.4132-08.2009

Wagner, M., Rihs, T. A., Mosimann, U. P., Fisch, H. U., & Schlaepfer, T. E. (2006). Repetitive transcranial magnetic stimulation of the dorsolateral prefrontal cortex affects divided attention immediately after cessation of stimulation. Journal of Psychiatric Research, 40(4), 315–321. 10.1016/j.jpsychires.2005.06.001

Wang, L., Li, X., Hsiao, S. S., Lenz, F. A., Bodner, M., Zhou, Y.-D., & Fuster, J. M. (2015). Differential roles of delay-period neural activity in the monkey dorsolateral prefrontal cortex in visual–haptic crossmodal working memory. Proceedings of the National Academy of Sciences, 112(2), E214–E219. 10.1073/pnas.1410130112

Watkins, A. J. (2005a). Perceptual compensation for effects of echo and of reverberation on speech identification. Acta Acustica United with Acustica, 91(5), 892–901. https://www.ingentaconnect.com/content/dav/aaua/2005/00000091/00000005/art00010

Watkins, A. J. (2005b). Perceptual compensation for effects of reverberation in speech identification. The Journal of the Acoustical Society of America, 118(1), 249–262. 10.1121/1.1923369

Watkins, A. J., & Makin, S. J. (2007). Steady-spectrum contexts and perceptual compensation for reverberation in speech identification. The Journal of the Acoustical Society of America, 121(1), 257–266. 10.1121/1.2387134

Watkins, P. V., & Barbour, D. L. (2008). Specialized neuronal adaptation for preserving input sensitivity. Nature Neuroscience, 11(11), 1259–1261. 10.1038/nn.2201

Weisser, A., Buchholz, J. M., Oreinos, C., Badajoz-Davila, J., Galloway, J., Beechey, T., & Keidser, G. (2019). The Ambisonic Recordings of Typical Environments (ARTE) database. Acta Acustica United with Acustica: The Journal of the European Acoustics Association (EEIG), 105(4), 695–713. 10.3813/aaa.919349

Wen, B., Wang, G. I., Dean, I., & Delgutte, B. (2009). Dynamic range adaptation to sound level statistics in the auditory nerve. The Journal of Neuroscience: The Official Journal of the Society for Neuroscience, 29(44), 13797–13808. 10.1523/JNEUROSCI.5610-08.2009

Wischnewski, M., & Schutter, D. J. L. G. (2015). Efficacy and Time Course of Theta Burst Stimulation in Healthy Humans. Brain Stimulation, 8(4), 685–692. 10.1016/j.brs.2015.03.004

Zahorik, P. (2009). Perceptually relevant parameters for virtual listening simulation of small room acoustics. The Journal of the Acoustical Society of America, 126(2), 776–791. 10.1121/1.3167842

Zahorik, P., & Brandewie, E. J. (2016). Speech intelligibility in rooms: Effect of prior listening exposure interacts with room acoustics. The Journal of the Acoustical Society of America, 140(1), 74–86. 10.1121/1.4954723

Zahorik, P., & Wightman, F. L. (2001). Loudness constancy with varying sound source distance. Nature Neuroscience, 4(1), 78–83. 10.1038/82931

Zekveld, A. A., Heslenfeld, D. J., Festen, J. M., & Schoonhoven, R. (2006). Top-down and bottom-up processes in speech comprehension. NeuroImage, 32(4), 1826–1836. 10.1016/j.neuroimage.2006.04.199

Zikopoulos, B., & Barbas, H. (2006). Prefrontal projections to the thalamic reticular nucleus form a unique circuit for attentional mechanisms. The Journal of Neuroscience: The Official Journal of the Society for Neuroscience, 26(28), 7348–7361. 10.1523/JNEUROSCI.5511-05.2006.

